# A reinforcement-based mechanism for discontinuous learning

**DOI:** 10.1101/2022.05.06.490910

**Authors:** Gautam Reddy

## Abstract

Problem-solving and reasoning involve mental exploration and navigation in sparse relational spaces. A physical analogue is spatial navigation in structured environments such as a network of burrows. Recent experiments with mice navigating a labyrinth show a sharp discontinuity during learning, corresponding to a distinct moment of ‘sudden insight’ when mice figure out long, direct paths to the goal. This discontinuity is seemingly at odds with reinforcement learning (RL), which involves a gradual build-up of a value signal during learning. Here, we show that biologically-plausible RL rules combined with persistent exploration generically exhibit discontinuous learning. In tree-like structured environments, positive feedback from learning on behavior generates a ‘reinforcement wave’ with a steep profile. The discontinuity occurs when the wave reaches the starting point. By examining the nonlinear dynamics of reinforcement propagation, we establish a quantitative relationship between the learning rule, the agent’s exploration biases and learning speed. Predictions explain existing data and motivate specific experiments to isolate the phenomenon. Additionally, we characterize the exact learning dynamics of various RL rules for a complex sequential task.

## INTRODUCTION

As we walk the streets of a city, we rapidly figure out paths to new spots after a few times visiting them. For nesting animals, foraging between new locations and their nests in structured environments is an essential aspect of their survival. Rats constantly navigate within a complex underground network of burrows to expand their stores of food [1]. Navigating from point A to point B in a structured space requires different strategies compared to a similar task on a flat, open field. In the latter, navigation often involves geometric calculations of distances and angles based on celestial cues, compasses or landmarks. In a burrow, on the other hand, a rat needs to learn which way to turn at each intersection and benefits from understanding the relationship between places within the network.

The relational structure of mazes offers a well-controlled experimental paradigm to identify biological algorithms for navigating structured environments. Early laboratory experiments on learning algorithms, and animal behavior at large, involved rats navigating a maze [2–7]. Rats rapidly learn to navigate to a rewarding location within the maze, which often develops into a habitual action sequence resistant to subsequent changes such as the addition of a shortcut. These experiments and others led to the hypothesis that learning entailed the fixation of stimulus-response relationships due to a reward [6–9]. A parallel set of experiments showed that the structure of the maze could be learned during exploration without any significant reward, termed as latent learning [10]. Latent learning presumably proceeds through the formation of a ‘cognitive map’, which can be flexibly re-used when the animal needs to generalize to a novel situation [11–13]. This dichotomy between behavioral stereotypy and flexibility is analogous to the modern dichotomy in computational reinforcement learning (RL) between direct and indirect learning, often implemented using model-free and model-based methods respectively [14–16]. However, the specific learning algorithms that animals use to navigate and the circumstances under which one system or the other is employed remain unclear.

Recent developments in deep-learning-based behavioral tracking methods [17–19] allow for following mice in labyrinthine mazes for extended periods of time. In an elegant experiment [20], mice were allowed to navigate (in the dark) an unfamiliar maze structured as a depth-six binary tree (Figure 1a). In each experiment, a mouse moves freely between a cage (marked as home in Figure 1a) and the maze. Markerless pose estimation [17] is used to track its movements continuously over seven hours. Ten of the twenty mice were water-deprived and a water reward was renewed every 90 seconds from a port at one end of the maze (marked as a water droplet in Figure 1a). Results recapitulate aforementioned studies: mice exhibit rapid learning and eventually execute a quick action sequence from home to the water port. In addition, mice persistently explore the maze with exploration biases which are remarkably consistent across rewarded and unrewarded animals.

**FIG. 1.**
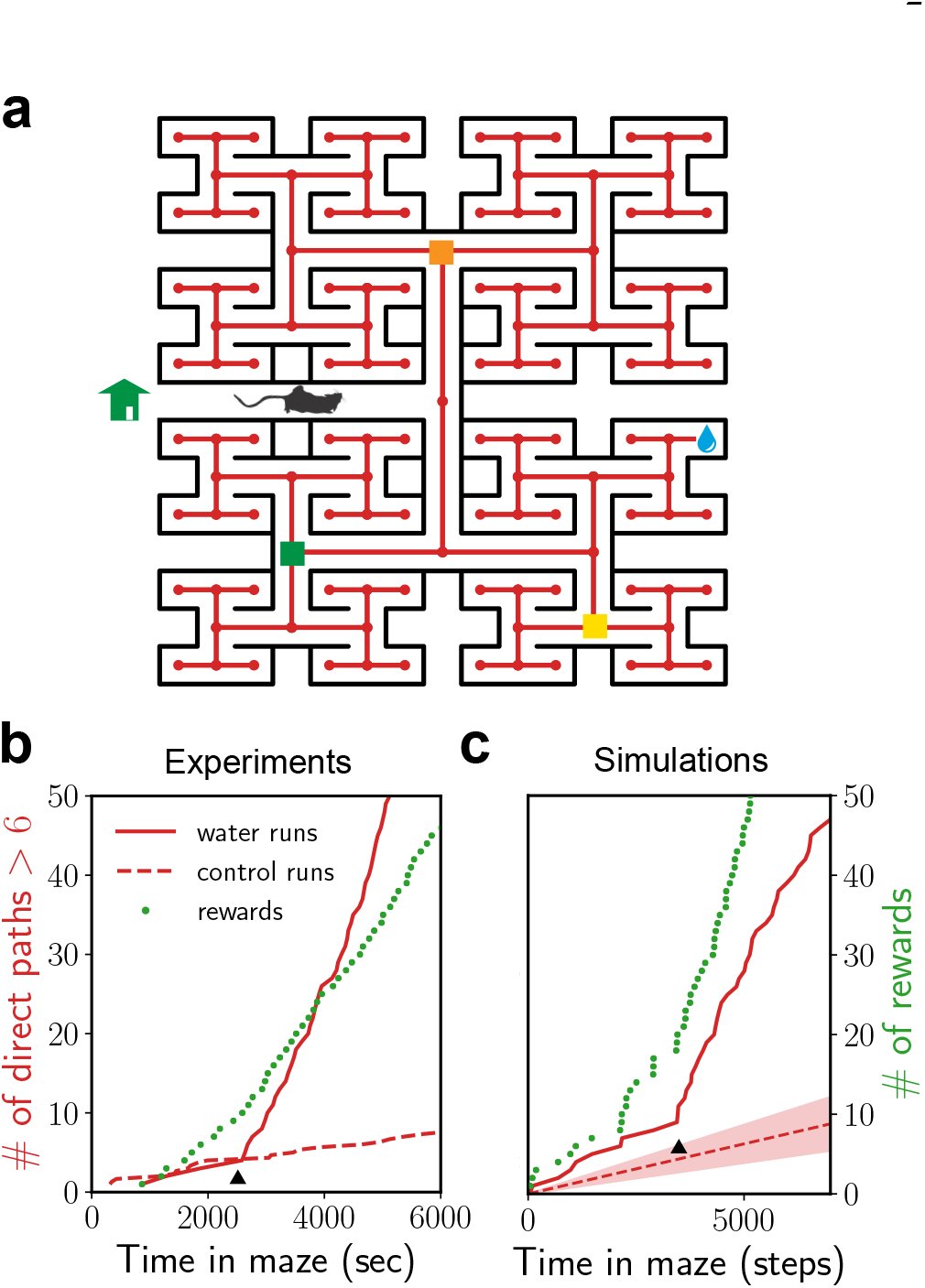
Discontinuous learning curves in mice experiments and RL simulations. (a) A schematic of the depth-6 binary tree maze used in experiments [20] and RL simulations. In each episode of the simulation, the agent begins at home and navigates the directed graph delineated by the maze (red) until it finds the reward. Three intersections (orange, green, yellow) that the mice have to pass through when executing a direct path of length > 6 are marked. (b) The cumulative number of direct paths of length > 6 (red) and acquired rewards (green) from an individual mouse. The rate of direct paths shows a discontinuity at a distinctive moment (black arrow). The dashed red line corresponds to length > 6 direct paths to control nodes. (c) Same as in (b) for RL simulations. See Figure S2 for more examples.

Intriguingly, the probability that mice take a direct path of > 6 correct binary choices towards the water port exhibits a sharp discontinuity, similar to an ‘a-ha’ moment of sudden insight (Figure 1b), and persists for the rest of the experiment. This moment can occur well after the animal acquires reward for the first time, which distinguishes this phenomenon from one-shot learning. Discontinuous learning curves have also been measured in a variety of other behavioral experiments [21]. RL algorithms reinforce correct actions in increments through accumulated experience. This intuition would suggest that RL-based learning is presumably incompatible with step-like learning curves. The availability of the full history of decisions made by mice within the maze presents a unique opportunity to identify the mechanism behind step-like learning curves.

In this manuscript, we use numerics and analytical calculations to rationalize the empirically observed discontinuous learning curves, and identify environmental architectures where we should generically expect such discontinuities. We present four main contributions. First, we use inverse reinforcement learning to decouple and analyze the influence of reward-based learning on the exploratory behavior measured in ref. [20]. In this setting, we show using agent-based simulations that persistent exploration combined with simple RL rules reproduce discontinuous learning. Second, we develop a general framework for RL-based sequence learning on tree-structured relational graphs. We use this framework to explain why RL algorithms will generically lead to discontinuous learning curves in such structured environments. Third, we develop a nonlinear, continuous-time model, which accurately captures the dynamics of reinforcement propagation in different exploration regimes. This model extends to commonly used model-free and model-based variants of RL, whose dynamics are analytically quantified. Finally, a re-analysis of experimental data lends further support for the theory and motivates specific experiments to isolate the phenomenon.

## RESULTS

### Discontinuous learning in RL simulations

We begin by specifying an RL model closely following the experimental setup of [20] (Figure 1a). The model is defined by the states (*s*), how the state changes when a certain action (*a*) is taken and the expected reward for each state-action pair, *r*(*s, a*). The states determine the information the agent can use to make a decision. Consistent with the history-dependence of exploratory behavior measured in experiments, we assume the agent knows which specific intersection it is currently at and where it is coming from. That is, the states are the directed edges of the graph that delineates the maze in Figure 1a. When the agent arrives at an intersection along a certain corridor, it has three choices: it can either choose to go along any of the corridors at that intersection or back where it came from. A fixed reward (*r*) is delivered in the corridor leading to water.

Upon finding the reward, the agent is reset at the starting point (marked in Figure 1a) and the simulation is repeated. This episodic formulation departs from the experimental setting; we find that an agent placed in an environment with delayed reward renewal (as in the experiment) often learns a degenerate policy which oscillates back and forth at the water port for the rest of the simulation. Of course, a mouse recognizes that water does not immediately reappear after it has been consumed (even if it does not know the precise renewal time) and explores the maze before eventually returning to the water port. For simplicity, we have used an episodic formulation instead of explicitly modeling this time delay.

An RL model is specified by the policy and the learning rule. We use a modified version of the standard softmax policy [14], which chooses actions with a log-probability proportional to their expected long-term reward or value, *q*(*s, a*), of taking action *a* at state *s*. Specifically, actions are chosen randomly with probability 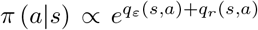 upto a normalization constant. Here, we have split *q*(*s, a*) into two terms, *q_ε_*(*s, a*) and *q_r_*(*s, a*). *q_ε_* is the *intrinsic* value the agent receives on taking an action at that state and is kept fixed throughout learning. *q_r_* is the *extrinsic* value, which is initially set to zero and is modulated by reward-based learning. Before learning, the agent makes stochastic exploratory choices based on *q_ε_*(*s, a*), which is presumably set by an innate bias or guided by knowledge external to the present task. This term is included in our RL model to explain the observed exploratory behavior of unrewarded and rewarded mice. As learning progresses, these exploratory choices are influenced by the reward, which biases the agent towards rewarding actions (or avoids costly ones). The randomness of the policy is set by the magnitudes of *q_ε_* and *q_r_*, whereas the influence of the reward on exploration is set by their ratio.

This split between intrinsic and extrinsic rewards allows us to examine *in silico* the influence of a learning rule on natural behavior. We first determine *q_ε_* from the behavior of unrewarded mice in experiments using maximum entropy inverse reinforcement learning (MaxEnt IRL [22, 23], Figure S1a). MaxEnt IRL finds the maximum entropy policy and the associated reward function that best explain observed behavioral trajectories (see Methods and SI for a brief overview of MaxEnt IRL). Next, we enable learning by specifying a biologically-plausible temporal-differences learning rule [14, 24–27]. Specifically, *q_r_* is updated using the learning rule:

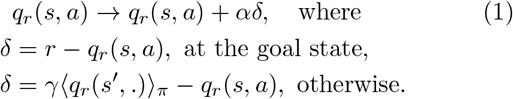

*δ* is the reward prediction error and the expectation above is with respect to the policy the agent uses at the next state (*s*′). The discount factor *γ*, which takes values between 0 and 1, is commonly used to introduce an effective time horizon and regularize the value function. Since our stochastic policy implicitly regularizes the value, we set *γ* = 1 throughout this paper. By comparing the best fit *q* values obtained from MaxEnt IRL for rewarded and unrewarded mice, we estimate the reward as *r* ≈ 2 (Figure S1c,d). The remaining free parameter, *α*, scales the rate of learning. Similar to the learning curves from experiments shown in Figure 1b we track the cumulative number of rewards acquired by the agent and the cumulative number of long direct paths (length > 6) to the goal from distant locations in the maze.

Simulated RL agents exhibit rapid learning similar to those observed in experiments. Importantly, the rate of taking a long direct path deviates discontinuously from the default rate (i.e., as expected from pure exploration) at a distinctive moment during learning, reproducing the ‘sudden insight’ phenomenon observed in experiments (Figure 1c). This phenomenon is reproduced during reruns with variability comparable to the variability observed across mice in experiments (Figure S2a). Fitting the rate of direct paths using a logistic function, we find that the transition can be localized to within fewer than three trials in about half of the runs (Figure S2b).

### Goal-oriented navigation on tree-like relational graphs

To identify the mechanism that underpins the sharp transition in learning, we now develop a framework for goal-oriented navigation on tree-like relational graphs. We use this framework to reproduce the discontinuous learning phenomenon, develop a mathematical theory that captures the learning dynamics and highlight the essential ingredients that lead to the phenomenon.

In this task, the agent traverses a relational graph (a directed graph whose edge labels specify the action or relationship between two states) from a fixed starting point to a goal where it receives a reward (Figure 2a). We track its progress in finding the direct path (highlighted in Figure 2a) by accumulating experience across multiple episodes. We wish to consider graphs that capture the core features of a structured environment such as roads on a university campus or abstract knowledge graphs [28]. Specifically, we require: 1) discrete decision points and choices, 2) the graph is sparse, namely, the number of paths of comparable length to the direct path is small (unlike a Manhattan-like grid), and 3) long, branching side paths which lead to dead-ends.

**FIG. 2.**
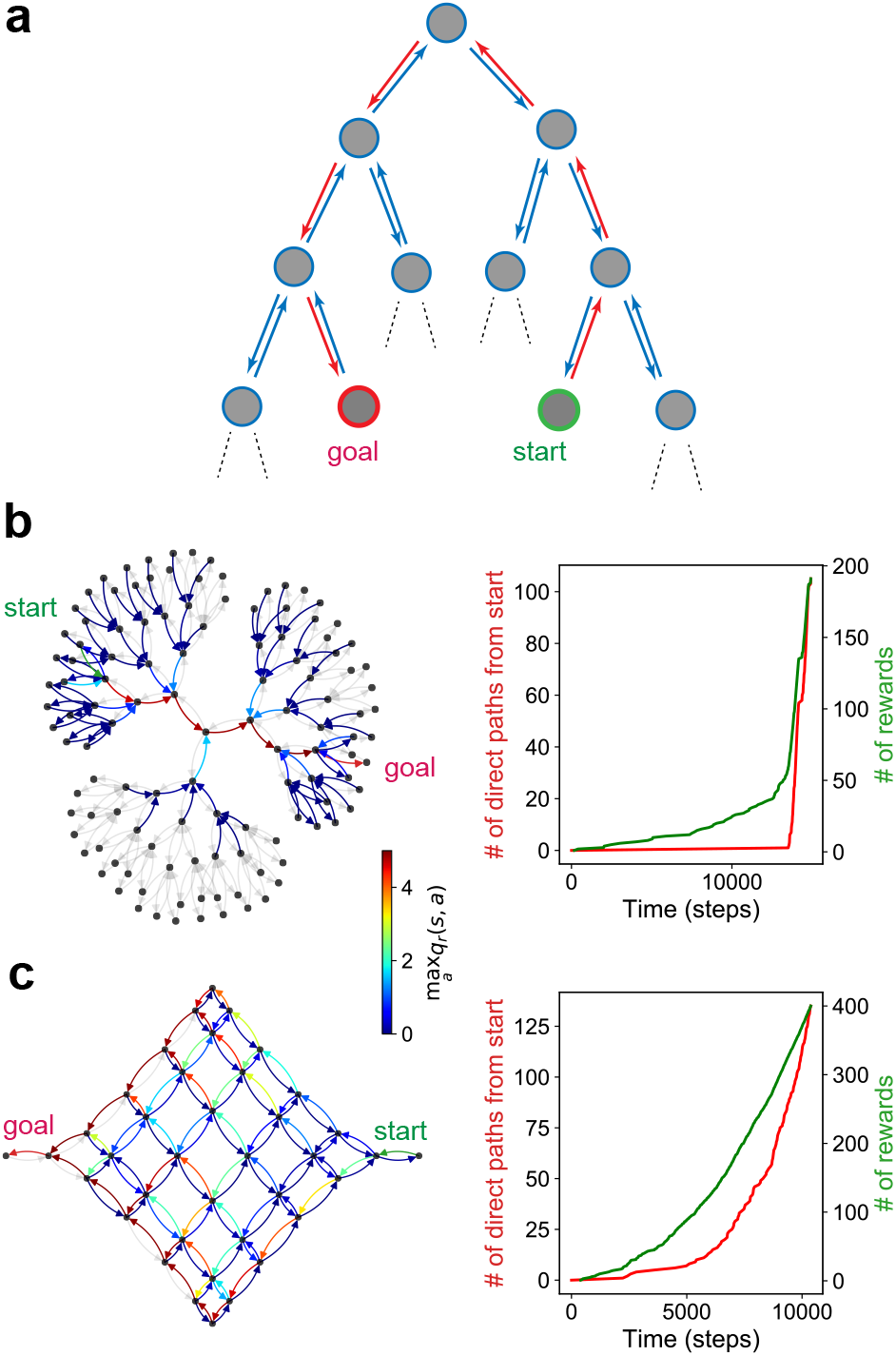
Reinforcement waves during sequence learning on relational graphs. (a) We consider a task where the agent traverses the directed edges of a relational graph to navigate from start (green) to goal (red). The direct path from start to goal is highlighted in red. (b,c) A discontinuous learning curve for a balanced ternary tree and a smooth learning curve for a Manhattan-like 6 × 6 grid. In all simulations, we use a standard softmax policy (*q_ε_* = 0) with *α* = 0.1, *r* = 5. The colors show the *q_r_* value of the best action at each state (directed edge). The gray edges have maximum *q_r_* value less than 10^−3^.

A large class of graphs that satisfy the above three requirements and yet sufficiently simple to allow for an in-depth quantitative analysis are tree-structured graphs (Figure 2a), which include the maze architecture from the experiments. Simulating an RL agent in a balanced ternary tree (Figure 2b), we find a sharp discontinuity in the rate of taking the direct path from the start to the goal. Examining the dynamics of reinforcement propagation shows that the reinforcement signal primarily propagates along the direct path (Figure 2b) and that the discontinuity occurs precisely when the reinforcement signal reaches the start (Movie S1). In contrast, an RL agent in a Manhattan-like 6 × 6 grid leads to diffuse propagation of the reinforcement signal and a smooth learning curve (Figure 2c, Movie S2). In Figures S3 and Movies S3, S4 we present the learning curves for four additional architectures: a binary tree where the length of the corridors agent is explicitly modeled, a binary tree where the agent is allowed to reverse its direction and two random graphs with different sparsities. We observe discontinuous learning curves for all of these architectures except for the dense random graph, highlighting that the task structure plays a role in whether discontinuous learning curves are observed.

The structure of tree-like graphs enables us to identify elements of the graph topology and learning dynamics that lead to discontinuous learning. The key insight is that the full complexity of sequence learning on a tree-like graph can be reduced to analyzing the learning dynamics on a simpler linear track with side paths represented as single nodes, as shown in Figure 3a. Specifically, recall that for tree-like graphs, the side paths necessarily lead to dead-ends. On encountering a dead-end, the agent will turn back and eventually re-encounter the direct path. The agent’s movements in a side path can thus be represented as a single node noting that if the agent goes in, it will surely return back. When the agent returns back from the side path, it can either choose to go towards or away from the goal.

**FIG. 3.**
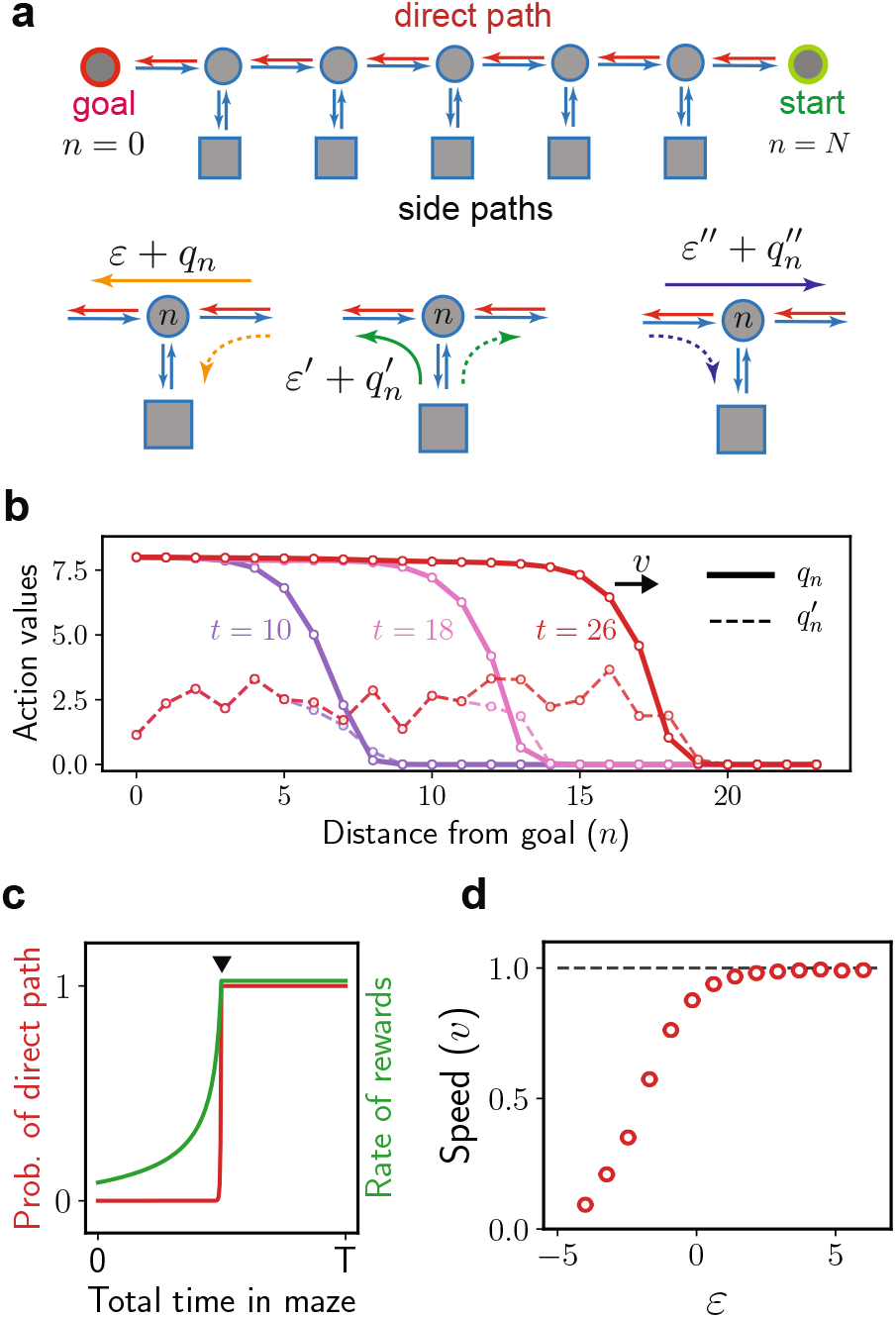
Wave-like reinforcement propagation during sequence learning on treelike graphs. (a) An equivalent representation of the tree-structured graph in Figure 2a highlighting the direct path and the possible branches into side paths at each intersection along the direct path. Note that while each side path is shown as a single node with reflecting boundaries, these represent long detours which will lead to a dead-end, forcing the agent to turn back and eventually return to the direct path. The exploration biases *ε, ε′, ε′′* and the corresponding reward-modulated biases 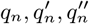 for the three cases of going towards the goal on the direct path (left), towards the goal from the side path (middle) and away from goal on the direct path (right) are shown. (b) The learned values *q_n_* and 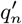 for three snapshots showing the propagation of the rein-forcement wave. (c) An illustration showing the discontinuity in the probability of a direct path and the rate of rewards. The discontinuity occurs the moment the wave hits the starting point, see Movie S5. (d) The speed of the wave for a range of *ε*. Smaller *ε* values correspond to more difficult tasks.

We emphasize two points that allow this simplification. First, even though we have used a single node with reflecting boundaries to represent the side paths (Figure 3a), an agent may spend a considerable amount of time exploring each of these side paths. Since the time spent within the side path does not influence reinforcement propagation on the direct path, we can safely assume the agent spends a single step on the side path. Note that the discontinuity in learning is sharper if we suppose the agent spends longer than a single step in each side path. Second, as long as the side path is sufficiently long, it is unlikely that the reinforcement signal will propagate through the entire side path and bias the agent to go *into* the side path. Therefore, we may ignore the details of the dynamics within the side path and assume that the *q_r_* value of going into the side path remains at zero. It is important to note that the agent may still learn to turn towards the goal when *exiting* a side path.

The agent’s exploration biases (specified by *q_ε_*) play an important role in determining the qualitative character of the learning dynamics. A key parameter is the probability of continuing towards the goal along the direct path whose corresponding *q_ε_* value we denote *ε* (Figure 2b). We have assumed a homogeneous *ε* for simplicity. The discontinuous learning phenomenon is still observed if this assumption is relaxed (see for example Figure 1c where the empirically derived *q_ε_* values are heterogeneous). By varying *ε*, we examine how the agent’s initial exploration and learning dynamics depends on the agent’s bias towards taking the correct actions. When *e^ε^* ≫ 1, the agent continues on the direct path for long stretches and rapidly reaches the goal. In this trivial case, the graph effectively reduces to a linear track without side paths that stretches from the starting point to the goal. In the opposite limit, *e*^−*ε*^ ≫ 1, correct actions along the direct path are rare. To make progress, the agent would have to take constant detours towards the goal through side paths, whose probability is set by the corresponding value *q_ε_* = *ε*′ (Figure 3a). Clearly, if the probability of going towards the goal both along the direct path and through side paths is small (*e*^−*ε′*^, *e*^−*ε*^ ≫ 1), the agent is very unlikely to make it to the goal. Thus, whether the agent makes any learning progress whatsoever will depend on the exploration biases. We find that for large graphs, the exploration statistics display three sharply delineated regimes depending on the net probability of going towards the goal *vs* back towards the start (SI). If this net probability is negative, the ‘cautious’ agent constantly returns to the starting point and does not learn the task. When the net probability is positive, the ‘adventurous’ agent on average ventures closer to the goal. The marginal case of zero net probability leads to diffusive exploration.

### The mechanistic basis of discontinuous learning curves

We now examine the learning dynamics generated by the rule, (1), beginning with RL simulations on the reduced architecture shown in Figure 3a followed by a theoretical analysis. Since actions that lead the agent away from the goal are never reinforced during learning, only the *q_r_* values for continuing along the direct path towards the goal (*q_n_*), and turning towards the goal when exiting the side path 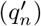 at each intersection *n* should be tracked (we use *n* = 0 and *n* = *N* for the goal and start respectively, see Figure 2b). Figure 3b shows *q_n_* and 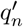 at three time points (in units of 1*/α* episodes), highlighting the wave-like propagation of the value, *q_n_* (Movie S5). The learning curves show a sharp discontinuity (Movie S5, Figure 3c), which occurs precisely when this wave reaches the starting point. Total learning time is determined by the wave’s speed, which we measure as the number of intersections on the direct path the wave crosses every 1*/α* episodes. Tracking the half-maximum of *q_n_*, we find that the wave travels at a constant speed, *v* (Movie S5). Simulations across a range of *ε* show the speed saturating at *v* = 1 for *ε* ≳ 1, which decreases to zero with decreasing *ε* (Figure 3d), hinting at distinct regimes. The factors that determine the speed and profile of the wave will be discussed in the following section.

The origin of discontinuous learning and ‘reinforcement waves’ can be intuitively understood by examining how learning operates at each intersection. We highlight three factors: 1) the correct action at an intersection is only reinforced if the action at the subsequent intersection is reinforced, implying that the chain of reinforcement has to travel backwards from the goal, 2) when an intersection is sufficiently reinforced, the probability of the correct action at that intersection increases by a large factor as long as the reward is sufficiently large (*e*^*r*+*ε*^ ≫ 1). Since the rate of traveling directly from start to goal is the product of the probabilities of taking the correct action at each intersection, this rate will increase rapidly when the wave reaches the start, and 3) if the agent is unlikely to take the correct action at a certain intersection (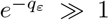 for that action), reinforcement is applied through a few rare events until the intrinsic bias is overcome, *q_r_* + *q_ε_* > 0. Since the probability of taking the correct action in turn increases rapidly with reinforcement, the learning curve for taking the correct action at *each* intersection will appear step-like.

The first factor emphasizes why we should expect the reinforcement signal to propagate backwards from the goal to the starting point. The second factor highlights the fact that the observable (i.e., the probability of taking the direct path) is a steep, nonlinear function of the underlying dynamical variables. The third point explains why the wavefront has a steep profile (Figure 3c). Put together, these three factors imply that when the task is non-trivial, the wave of reinforcement marches backward from the goal, reinforcing correct actions one intersection at a time with step-like learning at each intersection. The observed discontinuous transition in learning occurs when the wave reaches the starting point.

### A nonlinear, continuous-time model accurately captures the dynamics of reinforcement propagation

This intuitive picture can be made mathematically precise by examining the effects of the learning rule, (1), on *q_n_* and 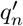. We summarize the results here and refer to the SI for full details. When *α* ≪ 1, we find that their expected change, 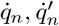, over 1*/α* episodes is given by

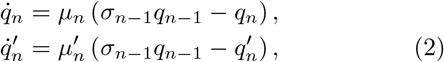

where 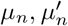 are the average number of times per episode the agent crosses intersection *n* through the direct path or the side path respectively, and *σ_n_* is the probability of continuing along the direct path at intersection *n*. In general, *μ_n_* and 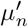 depend on the transition probabilities and thus the values at every intersection in the graph. The analysis is made tractable by noticing, first, that the ratio 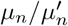 is determined by the relative probability of taking the correct action at intersection *n* through the direct path *vs* the side path. Second, no learning occurs outside of the front and bulk of the wave. Lastly, learning at the front of the wave only happens when subsequent intersections are already sufficiently reinforced, which implies that the agent is likely to go directly to the goal immediately after crossing the front. Thus, in each episode, the intersection at the wave’s front is crossed just once on average, 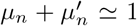. This relation combined with the expression for 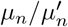 fix 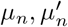. The 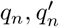’s obtained from numerical integration of (2) are in excellent agreement with the ones from full-scale RL simulations (Figure 4a). An analysis of (2) reveals two qualitatively distinct regimes of wave propagation with *e^ε^* ≫ 1 and *e*^−*ε*^ ≫ 1 as their asymptotic limits. We term these the expanding and marching regimes respectively. Maze architectures that could exhibit these two regimes are illustrated in Figure 4b,c.

**FIG. 4.**
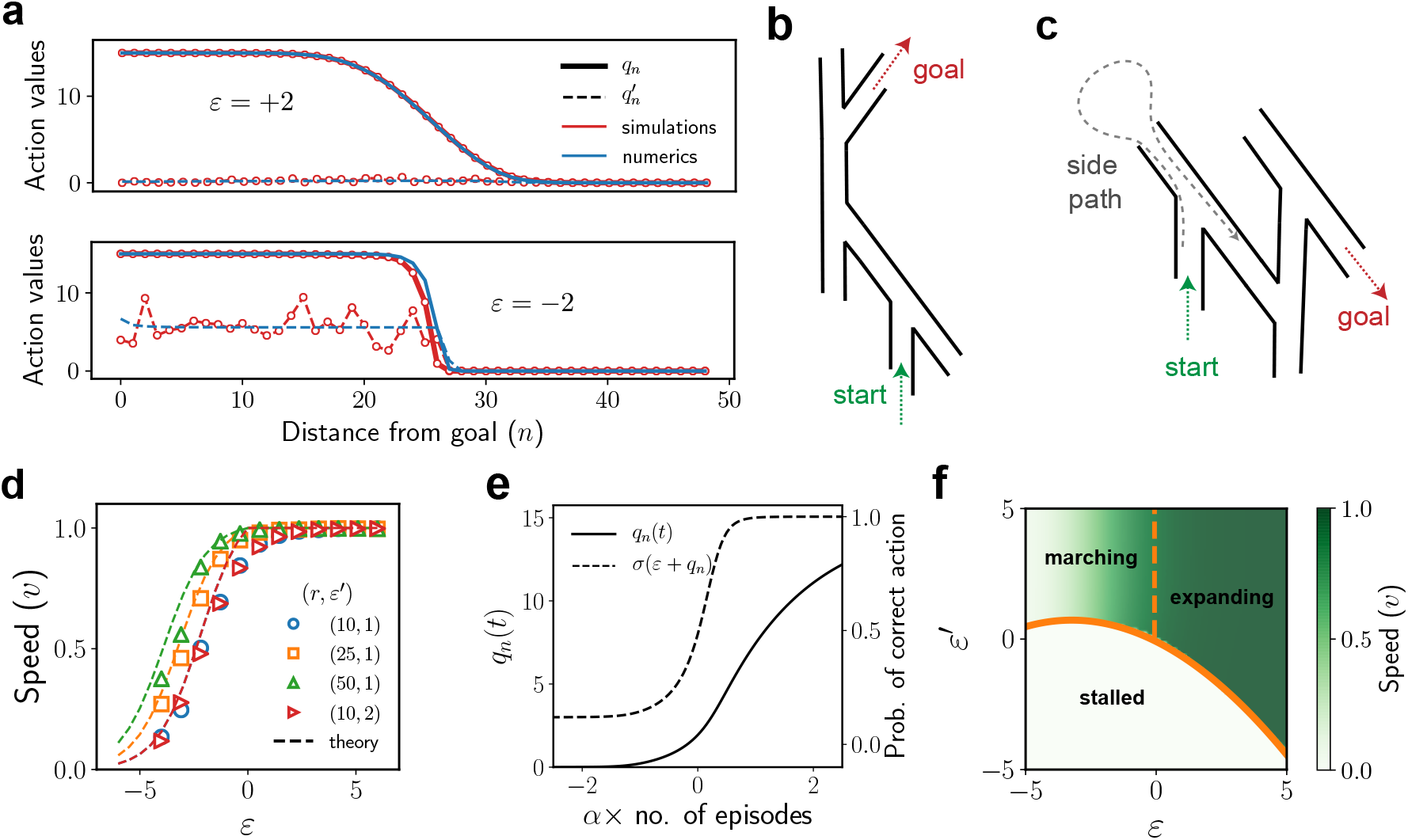
The expanding and marching regimes of wave propagation: numerics and theory (a) A snapshot of *q_n_* and 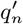 for *ε* = ±2, shown in red and blue from RL simulations and from numerically integrating (2) respectively. (b,c) Illustrations of how mazes with *e^ε^* ≫ 1 (b) and *e*^−*ε*^ ≫ 1 (c) can be constructed. (d) The theoretical prediction for the speed (dashed lines) closely aligns with the speed measured in the full RL simulations. The red and blue dashed lines are aligned. (e) The change in *q_n_*(*t*) when the wave passes through intersection *n*, shown here for *ε* = −2. The *x*-axis is centered at the moment when the probability of taking the correct action, *σ*(*ε* + *q_n_*) = 1/2. Note that learning at this intersection is localized to ≲ 1*/α* episodes. (f) The distinct learning regimes for a range of exploration parameters. Here *N* = 20, *ε*′′ = 0. We use *r* = 15 for panels a,e and *r* = 12 for panel f.

The expanding regime (*e^ε^* ≫ 1) corresponds to the trivial case where the agent is likely to traverse straight from the starting point to the goal. (2) leads to linear dynamics in this regime, which can be solved exactly. We find *q_n_*(*t*) = *rP* (*n, t*), where *P* (*n, t*) is the regularized lower incomplete gamma function. For large *n*, the half-maximum is at *n*_1/2_ = *t*, which explains the speed *v* = 1 observed in simulations for *ε* ≳ 1, and the width of the profile expands with time as 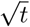.

In the marching regime (*e*^−*ε*^ ≫ 1), the negative *ε* leads to qualitatively different, non-trivial dynamics. Any step on the direct path that has previously been reinforced beyond |*ε*| is more likely to be traversed. When the reinforcement wave reaches an intersection *p* on the direct path that is yet to be reinforced to |*ε*|, the reinforcement of that step occurs through rare events until *q_p_* ⋍ |*ε*|. Meanwhile, the direct path for *n* < *p* is rapidly reinforced. The rare events at *p* combined with rapid reinforcement for *n* < *p* lead to a bottleneck at *p* and a steep wave profile. Once *q_p_* reaches |*ε*|, it is subsequently reinforced rapidly and *q*_*p*+1_ in turn begins to be slowly reinforced through rare events. Thus, the wave ‘marches’ forward reinforcing one step at a time. Computing the duration *τ* it takes to march one step will let us estimate the speed of the wave, *v* = *τ*^−1^.

The duration *τ* can be calculated by examining the nonlinear dynamics in the front (*n* = *p*) and bulk (*n* < *p*) of the wave (SI). The full dynamics in the bulk plays a role as the reinforcement received at the intersection *n* = *p* depends on the temporal dynamics of *q*_*p*−1_, which in turn depends on *q*_*p*−2_, and so on. However, it can be shown that the dynamics in the bulk are linear and exhibit self-similarity with period *τ*. Exploiting a conservation equation that results from these properties, we compute the wave speed as

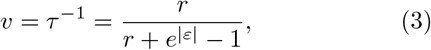

which is in excellent agreement with the speed measured in RL simulations (Figure 4d). The wave profile in the bulk is given by *q*_*n*−1_(*t*) = *r* − *β*(*r* − *q_n_*(*t*)), where *β* = −*τ*^−1^*W* (−*τe*^−*τ*^) and *W* (*x*) is the Lambert *W* function. Most of the learning at a certain intersection occurs in ≲ 1*/α* episodes (Figure 4e). Since the wave speed is less than one in the marching regime, each intersection is almost fully reinforced before the wave marches to the next one, thus quantifying the aforementioned intuitive argument that a step-like learning curve is observed at each intersection.

The results are summarized in Figure 4f, which depicts the expanding and marching regimes in addition to the ‘stalled’ regime corresponding to the exploration parameters where learning is largely absent.

### Other learning rules lead to reinforcement waves with altered speeds and profiles

Common variants of the *SARSA* rule [14] in (1) also lead to discontinuous learning via reinforcement waves, highlighting the generality of the phenomenon. A detailed analysis of each of these variants is presented in the SI, which we summarize here.

We find Watkins’ *Q*-learning, which uses a slightly modified version of the rule (1), leads to largely similar wave speeds and profiles. The advantage of *Q*-learning is that the *q_r_* values can be learned off-policy, i.e., the agent’s behavior is not necessarily derived from the learned *q_r_* values. To decouple the influence of learning on behavior, we use *Q*-learning together with an explorative agent that disregards the learned *q_r_* values. We find expanding waves irrespective of the exploration bias, suggesting that expanding waves are the ‘default’ dynamics without feedback in the structured environments considered here. Feedback due to learning leads to traveling waves with steeper profiles as observed in the marching regime. Both *Q*-learning and *SARSA* learn values from local updates, which constrains the wave speed to be at most one.

An alternative class of models build a model of the environment from experience, similar to a cognitive map, and update the values offline by sampling from the model (planning). We consider *Dyna-Q*, which implements a simple version of this general idea. Specifically, *Dyna-Q* first learns a model of future states and rewards for every state-action pair it encounters during the task. At each step, it samples *n_p_* state-action-state-reward transitions from the model and updates their corresponding values. We show that *Dyna-Q* applied to our setting leads to the same behavior as (1) with an enhanced learning rate (1 + *n_p_*)*α*. Intuitively, when the agent plans, learning which otherwise occurs only through physical exploration is sped up due to mental exploration. However, since both physical and mental exploration employ the same search process, the result is a simple scaling of the learning rate.

Another common variant with non-local updates is *SARSA* combined with eligibility traces, which are an efficient, biologically-plausible mechanism for enhancing learning speed when rewards are sparse [14, 29]. Instead of updating the value of the current state-action pair, eligibility traces effectively use the current reward prediction error to also update the *k* most recent state-action pairs. The exact learning dynamics can be calculated (SI) and are qualitatively similar to the *SARSA* case. In the expanding regime, eligibility traces scale the wave speed by a factor 1+*k*. The speed in the marching regime has a non-trivial, sub-linear relationship with *k* (Figure S4b), which can be computed from the theory using a self-consistent equation (SI). Intuitively, the speed increases with *k* since the front of the wave receives reinforcement from the intersection 1 + *k* steps along the direct path, which has a larger value compared to the subsequent intersection. In the limit *k* → ∞, we show that the speed converges to a maximum *v*_∞_ = *r/*(|*ε*| + *e*^|*ε*|^ − 1).

The theoretical predictions for the various learning rules are verified in simulations (Figures S4,S5).

## EXPERIMENTAL TESTS

In addition to reproducing the discontinuous learning curves observed in experiments, the theory provides predictions which can be immediately tested by re-analyzing the data from ref. [20]. Specifically, note that the learning curves in Figure 1b correspond to the number of direct paths greater than a certain length, namely, six. If the discontinuity in the learning curves is due to a reinforcement wave, this discontinuity should occur at a later time for direct paths beginning from farther nodes. This prediction should be contrasted with an alternative mechanism where sudden insight corresponds to the singular moment when the mouse has figured out the global structure of the environment and uses this knowledge to find direct paths from distant sections of the maze. The experimental data lends support for the former hypothesis, which show that the discontinuity is delayed for longer direct paths (Figure 5a). The time delay between these discontinuities provides an estimate of the wave speed. The smaller rate of taking direct paths for longer paths observed in Figure 5a can also be explained in our framework. The reward (estimated as *r* ≈ 2 previously) is not sufficiently large to fully overcome the stochastic, exploratory drive of the agent, leading to a significantly smaller probability of taking a longer direct path. This decreasing probability provides an estimate for the range of wave propagation, *N*_range_. The theory predicts that *N*_range_ and the speed of wave propagation should increase with increasing reward for *e*^−*ε*^ ≫ 1, which can be tested in future experiments. An intriguing possibility is to observe the transition in speed from the expanding to marching regimes by manipulating the exploration biases, for example by modifying the inclinations of the T-junctions in a complex maze (as illustrated in Figure 4b,c) or manipulating the number of branches at each intersection.

**FIG. 5.**
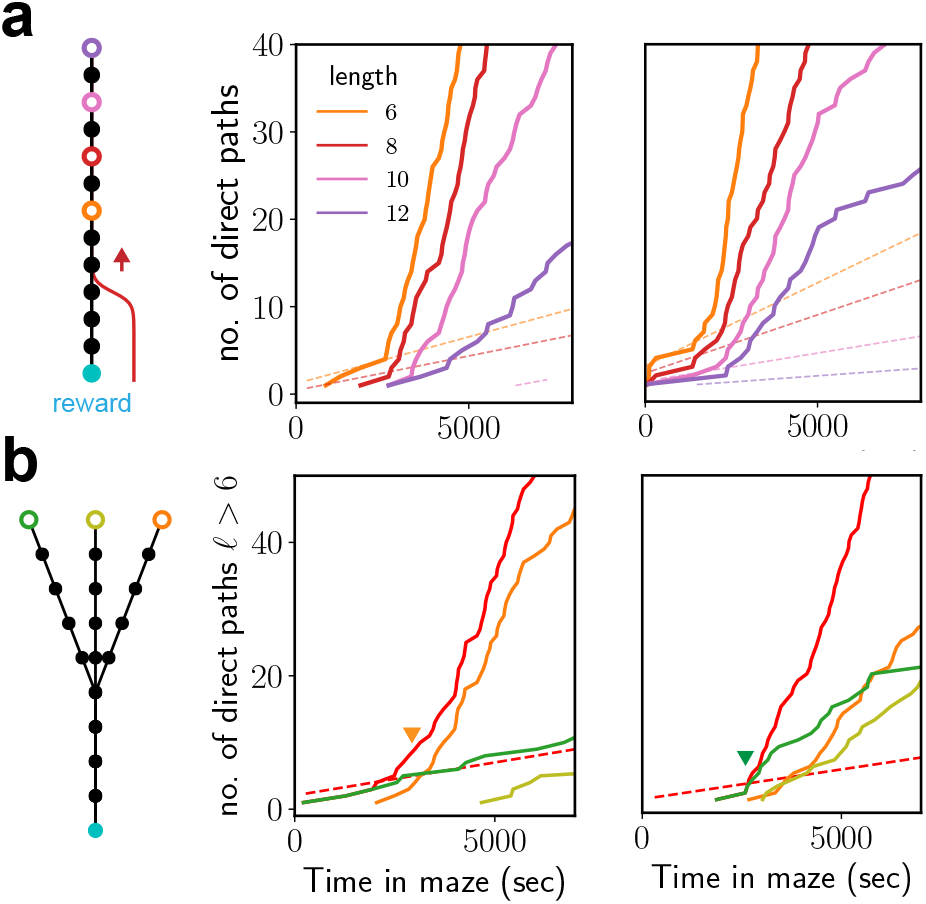
Wave propagation is consistent with experimental data. (a) Theory predicts that the reinforcement wave reaches locations further away from the goal at later times. Shown here are the cumulative number of direct paths of lengths at least 6, 8, 10 and 12 in orange, red, pink and purple respectively for two mice. The dashed lines are direct paths to control nodes. (b) Stochasticity in exploration and learning dynamics can lead to the wave reaching different intersections at different moments during learning. Shown here are direct paths of length > 6 from three distinct intersections in the maze (marked in Figure 1a with their respective colors) for two mice.

A potentially important confounding factor for observing a single, distinct discontinuity in the learning curves is when multiple paths of length comparable to the direct path are available. The speed at which the wave propagates along these competing paths depends on a number of factors, including their number, lengths and the exploration statistics within each path. If a competing path is fully reinforced earlier than the direct path, it can interfere with learning the direct path. Multiple paths can explain the variability observed in experimental trajectories. Indeed, the learning curves in Figure 1b,5 effectively average over all direct paths of certain lengths. If paths of similar lengths from distant nodes exhibit discontinuities with only slight delays, the averaged curve will appear smoother than when each path is observed separately. Consistent with this intuition, considering paths from specific locations in the experiment highlights the variability across mice in which of these paths contributes most to the discontinuity (Figure 5b).

Additional experiments designed similar to our setting in Figure 2a will provide crucial data to resolve sources of variability. Specifically, our analysis suggests examining direct paths between two specific start and goal locations in an episodic setting or equivalent. This will ensure that the measured learning curves do not reflect contributions from different locations in the maze, and highlight the passage of the wave along the direct path between these two nodes. Further, learning via reinforcement is not necessarily monotonic in the experimental setup of ref. [20], which makes it challenging to infer the progression of learning at each intersection directly from data. For example, if the animal samples the water port when reward is absent, the resulting reinforcement can be negative which leads to un-learning of the path towards reward. This non-monotonicity is absent in an episodic setting, and will lead to a clearer interpretation of the learning curves at each intersection.

## DISCUSSION

The discontinuous learning phenomenon observed in complex mazes and other learning tasks clashes with the intuition that RL-based algorithms make learning progress by incrementally reinforcing rewarding actions. Here, we have shown that a standard biologically-plausible RL rule consistently reproduces this phenomenon in simulations designed to reflect maze experiments and more generally during goal-oriented navigation in large, tree-like relational graphs. In such environments, the value signal propagates as a steep, traveling reinforcement wave, which sequentially reinforces correct actions along the path towards the goal. ‘Sudden insight’ occurs the moment the wave reinforces all the correct actions along the main path. Discontinuous learning curves arise due to a combination of the effectively one-dimensional task structure in tree-like structured environments, the local propagation of reinforcement and the positive feedback of reinforcement on behavior. These factors together with the agent’s innate exploration biases determine the dynamics of wave-like reinforcement propagation, including its speed and profile. The exploration biases play an important role as they determine if any learning occurs in the first place (the stalled regime), and if learning does progress, whether the learning dynamics are limited by the learning rule (expanding regime) or due to the low probability of taking the correct action (marching regime). While common model-free and model-based variants of the RL rule may enhance the learning speed and alter the wave’s profile, the qualitative characteristics of wave propagation are preserved.

Whether and under what contexts animals learn correct actions directly from experience or indirectly through a learned model of the environment is a long-standing debate. The ‘a-ha’ moment observed in the experiments of ref. [20] would naively appear to support the latter hypothesis. We have shown here that existing experimental data is consistent with the propagation of a reinforcement wave (Figure 5), and thus RL-based direct learning cannot be ruled out. Further experiments should reveal and verify the generality of the discontinuous learning phenomenon. The framework presented in this manuscript should help guide specific experiments to delineate direct and indirect learning (see Experimental Tests for further discussion).

We emphasize that the backward propagation of reinforcement arises as a straightforward consequence of the local RL rule applied in an environment where the goal state is the sole source of reward. However, as illustrated in Figure 2c and Figure S3d, not all graph architectures will display discontinuous learning under these RL rules. We have shown that the topology of large tree-like mazes (with appropriate exploration biases) supports discontinuous learning, but we expect to observe the phenomenon more generally if the graph satisfies certain notions of ‘sparsity’ (Figure S3c). This is because the sharp transition in learning is most salient in highly complex mazes where the direct path is non-trivial and paths other than the direct path are present but are poor solutions.

Competing paths lead to additional complexity, analogous to when a multitude of local minima compete with the global solution in non-convex optimization problems. Easily accessible competing paths which are of comparable length to the direct path may lead to non-trivial exclusion effects, effectively average out the learning curves and amplify variability due to minor differences in exploration biases across animals. Sudden, delayed improvements in generalization performance have been recently observed when neural networks are trained to solve small algorithmic tasks, a phenomenon that has been termed ‘grokking’ [30]. Preliminary theoretical work [31] suggests that the task structure imposes highly specific constraints on the representations that can achieve perfect generalization, and ‘sudden insight’ occurs when these constraints are fulfilled. This work and ours suggest that non-trivial constraints on good solutions imposed due to task structure might play an important role in the emergence of sudden learning phenomena.

Our analysis provides a complete characterization of the learning dynamics of various RL rules for a non-trivial sequential decision-making task, which is currently lacking. A key challenge in the theoretical analysis of RL algorithms is the feedback of learning on behavior, which makes the data distribution inherently non-stationary. In our setting, the non-stationarity is reflected by the dynamics of the wave during learning. We have shown that the front of the wave effectively acts as an absorbing boundary, which simplifies the analysis considerably. The learning speed is determined by the number of times the learning rule updates the value at the nose of the wave. Since this number itself depends on the value at the nose, the dynamics are nonlinear. In turn, since the value of the subsequent action depends on the value of the later actions within the bulk, the full interactions between the nose and the bulk of the wave will influence learning speed. We show that the learning speed cannot exceed a certain value due to the locality of the learning rule. Relaxing the locality constraint using eligibility traces enhances the learning speed by widening the value differential between the unreinforced action and the distal action from which it receives reinforcement. A model-based method which uses planning scales up the speed simply by scaling up the number of times it updates each action rather than due to a qualitative change in how reinforcement is propagated.

In specifying our model, we have made certain simplifications that do not capture the full complexity of animal learning. First, we have considered a discretized model of the state and action spaces. While this is a standard approximation, animals use continuous spatial representations and motor control. Standard computational RL rules, such as (1), have been fruitfully extended to deal with continuous state and action spaces, for instance, using function approximation and policy gradient methods [14]. Biologically plausible variants of these extensions have been proposed [32], including for mental exploration in simple mazes [33]. We expect the core intuition behind discontinuous learning to hold even for a more realistic model with continuous state and action spaces. A detailed analysis of RL dynamics for a model which takes these various factors into account is beyond the scope of current work (see Figure S3a, Movie S3 for preliminary results). Second, we have assumed that the animal has a unique representation of each corridor in the maze from the outset. Of course, this representation would have to be learned before the animal can assign and update the value of taking different actions at each corridor [34]. Our results should still apply if the timescale for “mapping” the environment is faster than the timescale for reinforcement learning. The hierarchical structure of neural network-based function approximators enables simultaneous learning of representations and values [35], but the timescales on which these processes operate in animals are unknown and presumably much shorter. An exciting future direction is to extend our framework to spatial navigation tasks with other graph topologies or when learning of proper actions is intertwined with the learning of continuous state representations.

## METHODS

### Extracting exploration statistics from data and hyperparameters for RL simulations

We use MaxEnt IRL (see SI for a brief introduction) to infer the exploration biases of unrewarded mice. As discussed in the Results section, the state space was chosen as the directed edges of the graph that delineate the maze in experiments, where the root of the tree corresponds to ‘home’. We pooled trajectories from all unrewarded mice, set *γ* = 0.8 and split the trajectories to length *T* = 12 (*T* should be at least the effective horizon ~ (1 − *γ*)^−1^ = 5 and choosing a large *T* slows inference). The choice of *γ* was motivated by the analysis in [20], which showed that a variable length Markovian model typically chooses ≲ 5 previous states to predict mice behavior. The *q_ε_* values are obtained from maximum likelihood estimation, specifically, from log *p*_**λ**,0_(*s, a*) after optimizing for ***λ*** (SI). Note that due to normalization the *q_ε_* values are determined only up to a constant additive term for each state.

To estimate *r*, we apply the above procedure to both unrewarded and rewarded mice. We calculate the difference between rewarded and unrewarded animals in the differences of the correct action’s *q* value and the effective *q* values of the other two incorrect ones (note 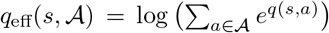). A subset of these values are shown in Figure S1, which shows that the correct actions leading to reward have a value differential of ≈ 2. Since the values of actions close to the reward after learning saturate at *r*, the value differential is an estimate of the reward, *r* ≈ 2. To ensure that this estimate is not significantly influenced by the habitual paths that go directly from home to goal, we repeat the above procedure excluding these paths (Figure S1). The estimate decreases slightly to *r* ≈ 1.5. In the RL simulations of the depth-6 binary tree maze, we use *r* = 2 and *α* = 0.33.

### Setup and notation for the RL framework for navigation on tree-structured graphs

A tree-structured graph can be cast as a linear track, as argued in the Results section and illustrated in Figure 2a,b. The linear track consists of *N* − 1 nodes on the direct path, *n* = 1, 2, …, *N* − 1. The agent starts each episode at node *n* = *N* and the reward is at the goal node *n* = 0. In addition to these nodes, the nodes from *n* = 1 to *N* − 1 each have a side path, which we label as 1_*b*_, 2_*b*_, …, (*N* − 1)_*b*_. The state space of the Markov decision process is the set of *directed edges* that connect the various nodes and the side paths as shown in Figure 2b. In other words, both the agent’s location in the graph and the direction in which it is headed matter. We denote (*n*_1_, *n*_2_) as the directed edge from *n*_1_ to *n*_2_.

The transition dynamics *P* (*s*′|*s, a*) are deterministic (note however that the policy *π*(*a*|*s*) is stochastic). At each directed edge, the agent can choose to go along the directed edges emanating from its current node, except for turning back, for e.g., the transition (*n* + 1*, n*) → (*n, n*+1) is disallowed. This simplifying assumption does not affect the results as the agent can effectively turn back by going into a side path and returning (*n* + 1*, n*) → (*n, n_b_*) → (*n_b_, n*) → (*n, n* + 1). The episode begins with the agent at the directed edge (*N, N* − 1). The directed edge pointing towards the goal node, (1, 0), is an absorbing state, i.e., the agent receives a reward *r* and the episode ends once the agent traverses that edge. We impose reflecting conditions at edges going into the side paths (*n, n_b_*) and the start node (*N* − 1, *N*).

The agent receives identical intrinsic exploration rewards at every intersection on the direct path. There are three directed edges leading to any node *n*, and we thus consider three cases at each node. These three cases are shown pictorially in Figure 2b. Since the agent can take two actions at each step and the policy only depends on differences of *q* values, we specify the *q* values for only one of the actions. The notation used for the three cases is introduced (see also Figure 2b).

1. the agent is on the direct path and going towards the goal, (*n* + 1*, n*): for the action corresponding to the agent continuing towards the goal (*n* + 1*, n*) → (*n, n* − 1), we denote *q_ε_* ≡ *ε, q_r_* ≡ *q_n_*.
2. the agent is on the side path *n_b_* and going towards *n*, (*n_b_, n*): for the action corresponding to the agent turning towards the goal (*n_b_, n*) → (*n, n* − 1), we denote *q_ε_* ≡ *ε*′, 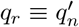.
3. the agent is on the direct path and going towards the start, (*n*−1, *n*): for the action corresponding to the agent continuing towards the start (*n* − 1*, n*) → (*n, n* + 1), we denote *q_ε_* ≡ *ε*′′, 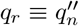.

The probabilities of taking the action described in each of three cases is denoted *σ_n_* ≡ *σ*(*ε* + *q_n_*), 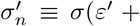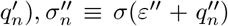, where *σ*(*x*) = 1/(1 + *e*^−*x*^) is the logistic function.

## ACKNOWLEDGMENTS

G.R thanks Andrew Murray and Venkatesh Murthy for useful comments on the manuscript. G.R was partially supported by the NSF-Simons Center for Mathematical & Statistical Analysis of Biology at Harvard (award number #1764269) and the Harvard Quantitative Biology Initiative.

## APPENDIX

## 1. Maximum entropy inverse reinforcement learning

In this Section, we present a brief introduction to Maximum Entropy Inverse Reinforcement Learning (MaxEnt IRL). See refs. [22, 23] and references therein for more details. MaxEnt IRL aims to find a reward function *r*(*s, a*) for each state-action pair (*s, a*) that guides a policy consistent with a set of *L* observed behavioral trajectories 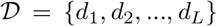. Each trajectory has *T* + 1 state-action pairs: 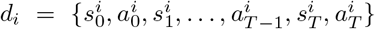. The transition matrix *P* (*s*′|*s,a*) is assumed to be known.

While the end goal is to estimate *r*(*s, a*), MaxEnt IRL proceeds by formulating an unsupervised learning task. Specifically, consider a generative model which assigns a probability *p*(*τ*) for trajectory *τ* such that the expected discounted reward, 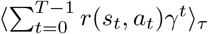 from the model matches the empirical discounted reward (note 0 ≤ *γ* < 1 is the standard RL discount factor). To do this, it is sufficient to ensure that the expected frequencies of all *s, a* pairs appropriately discounted match the empirical ones:

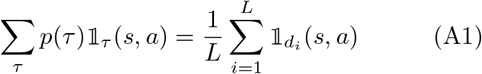

where the sum over *τ* is over all possible trajectories with *T* + 1 state-actions pairs, 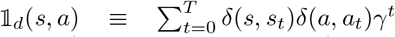 for a trajectory *d* = (*s*_0_, *a*_0_, *s*_1_, *a*_1_, …, *s_T_*, *a_T_*) and *δ* is the indicator function. Since multiple reward functions satisfy (A1), the inverse RL problem is ill-posed without additional constraints. MaxEnt IRL introduces an additional constraint by choosing the generative model *p*(*τ*) which satisfies (A1) *and* minimizes the relative entropy between *p*(*τ*) and a model *u*(*τ*) whose trajectories are generated by a random policy. In other words, MaxEnt IRL selects the “most random” policy which also satisfies (A1).

*p*(*τ*) is found by minimizing a variational objective with constraints imposed using Lagrange multipliers:

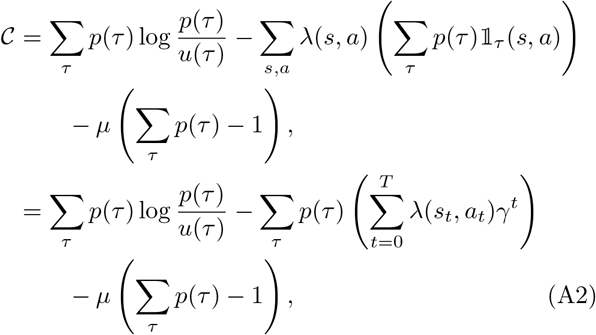

where *u*(*τ*) is defined below, the *λ*(*s, a*)’s are Lagrange multipliers which enforce (A1) and *μ* enforces normalization. Minimizing 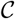 w.r.t *p*(*τ*) gives

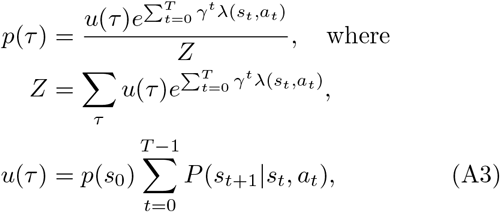

and *p*(*s*_0_) is the probability of the initial state. *u*(*τ*) is (up to a constant factor) the probability of trajectory *τ* under a policy which picks actions with equal probability. Observe that the discounted sum in the exponent of *p*(*τ*) is precisely the discounted long-term reward along *τ*. The Lagrange multipliers ***λ*** are thus interpreted as the rewards, *r*(*s, a*) → *λ*(*s, a*) for each *s, a*. *p*(*τ*) implicitly places an exponentially larger weight on policies that lead to rewarding trajectories relative to the uniform random policy corresponding to *u*.

The rewards ***λ*** are obtained by maximizing the log-likelihood 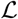 of the model over the sample trajectories:

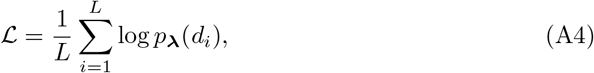

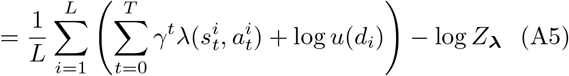

where the subscript ***λ*** is introduced to highlight the dependence on ***λ***. From (A3), we find

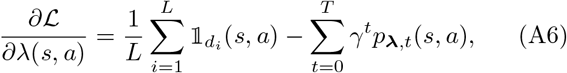

where *p*_***λ***,*t*_(*s, a*) is the marginal probability of encountering state-action pair *s, a* at time *t* w.r.t *p*. The Markovianity of the process enables efficient calculation of *p*_***λ***,*t*_(*s, a*) using a forward-backward algorithm (described below) similar to the one used to train Hidden Markov Models (HMM). When 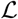 attains its optimum, we observe from (A6) that the constraint (A1) is automatically satisfied.

The forward-backward equations are similar to the HMM case except for an exponential weight:

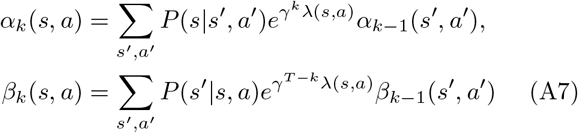

for *k* > 0 and *α*_0_ (*s, a*) = *p*_0_ (*s*)*e*^*λ*(*s,a*)^, 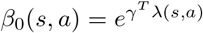. We have

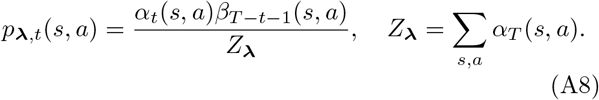

In practice, we compute the logarithm of the various quantities and use logΣ_*x*_ *e^x^* = max(*x*) + log(1 + Σ_*x*≠max(*x*)_ *e*^*x*−max(*x*)^) to prevent under-flow/overflow. The *λ*s are obtained using gradient ascent on 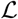 using standard optimization methods.

## 2. The dynamics of sequence learning on tree-structured relational graphs

In this Section, we expand on the dynamics of learning in the goal-oriented navigation task introduced in the main text. The dynamics are analyzed in full detail for the SARSA learning algorithm used in the main text and other commonly used RL rules. These alternative rules also display wave-like propagation of the value signal, with differences in the speed and front profiles for certain variants. Part of this Section is reproduced in the Methods, and is included here for completeness.

## Learning framework

An RL algorithm is defined by the learning rule and the (possibly non-stationary and stochastic) policy executed during learning. We consider a stochastic policy, where for each state-action pair (*s, a*), we have ln *π*(*a|s*) = *q*(*s, a*) − ln *Z*(*s*), where *q*(*s, a*) = *q_ε_*(*s, a*) + *q_r_*(*s, a*) and *Z*(*s*) is a normalization constant. *q_ε_*(*s, a*) represents fixed intrinsic rewards that drive exploration and determine the agent’s exploratory biases. The extrinsic rewards *q_r_*(*s, a*) (initially set to zero) are modulated by the learning rule. At the beginning of learning, the agent’s behavior depends solely on its exploratory biases. As learning proceeds and the agent acquires rewards, its behavior is modified by the increasing effect of the extrinsic rewards. We consider the learning dynamics for various learning rules discussed further below.

## Setup and notation

The linear track consists of *N* − 1 nodes on the direct path, *n* = 1, 2, …, *N* − 1. The agent starts each episode at node *n* = *N* and the reward is at the goal node *n* = 0. In addition to these nodes, the nodes from *n* = 1 to *N* − 1 each have a side path, which we label as 1_*b*_, 2_*b*_, …, (*N* − 1)_*b*_. The state space of the Markov decision process is the set of *directed edges* that connect the various nodes and the side paths as shown in Figure 2b. In other words, both the agent’s location in the graph and the direction in which it is headed matter. We denote (*n*_1_, *n*_2_) as the directed edge from *n*_1_ to *n*_2_.

The transition dynamics *P* (*s*′| *s, a*) are deterministic (note however that the policy *π*(*a*|*s*) is stochastic). At each directed edge, the agent can choose to go along the directed edges emanating from its current node, except for turning back, for e.g., the transition (*n* + 1*, n*) → (*n, n*+1) is disallowed. This simplifying assumption does not affect the results as the agent can effectively turn back by going into a side path and returning (*n* + 1*, n*) → (*n, n_b_*) → (*n_b_, n*) → (*n, n* + 1). The episode begins with the agent at the directed edge (*N, N* − 1). The directed edge pointing towards the goal node, (1, 0), is an absorbing state, i.e., the agent receives a reward *r* and the episode ends once the agent traverses that edge. We impose reflecting conditions at edges going into the side paths (*n, n_b_*) and the start node (*N* − 1*, N*).

1. the agent is on the direct path and going towards the goal, (*n* + 1*, n*): for the action corresponding to the agent continuing towards the goal (*n* + 1*, n*) → (*n, n* − 1), we denote *q_ε_* ≡ *ε, q_r_* ≡ *q_n_*.
2. the agent is on the side path *n_b_* and going towards *n*, (*n_b_, n*): for the action corresponding to the agent turning towards the goal (*n_b_, n*) → (*n, n* − 1), we denote *q_ε_* ≡ *ε*′, 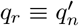.
3. the agent is on the direct path and going towards the start, (*n*−1, *n*): for the action corresponding to the agent continuing towards the start (*n* − 1*, n*) → (*n, n* + 1), we denote *q_ε_* ≡ *ε*′′, 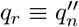.

The probabilities of taking the action described in each of three cases is denoted *σ_n_* ≡ *σ*(*ε* + *q_n_*), 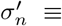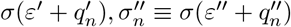, where *σ*(*x*) = 1/(1 + *e*^−*x*^) is the logistic function.

## Dynamics of exploration

We first consider the dynamics of exploration with no learning. An important byproduct of the calculation is the identification of the values of *ε, ε*′*, ε*′′ for which the agent finds the goal within a reasonable amount of time. These values delineate the parameter regime where learning occurs (the marching and expanding regimes in Figure 3c) *vs* the regime in which no learning occurs as the agent rarely finds the goal in the asymptotic limit *N* ≫ 1 (the stalled regime in Figure 3c). The upshot is that in these two cases the agent on average either drifts linearly towards the goal or constantly reverts back to the start. The marginal case corresponds to diffusive behavior.

To identify these regimes, we calculate the expected time for the agent, starting at (*N* − 1*, N*), to find the goal state. We additionally calculate the expected number of times the agent visits each state which is used further below for the analysis of learning dynamics. The calculation of both of these quantities is simplified by considering the dynamics of a simpler, equivalent Markov chain. Specifically, we observe that when the agent enters a node *n*, say along (*n* + 1, *n*), regardless of whether it goes into the side path or not, it either continues along its path, (*n, n* − 1), or turns back (*n, n* + 1). The probability of the former possibility, *k*_+_, is the sum of the probability that it moves to (*n, n* − 1) along the direct path, *σ*(*ε*), and the probability that it takes a detour through the side path, (1 − *σ*(*ε*))*σ*(*ε*′). The probability of turning back is 1 − *k*_+_. Similarly, the probability of continuing towards the start state when the agent is at (*n* − 1*, n*) is *k*_−_ = *σ*(*ε*′′) + (1 − *σ*(*ε*′′))(1 − *σ*(*ε*′)), and the probability of turning back to (*n, n* − 1) is 1 − *k*_−_. In summary, the probabilities that the agent continues towards the goal and towards the start are

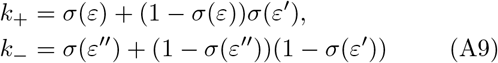

respectively. The dynamics of the Markov chain defined by these two parameters are easily computed.

## (a) Expected time to the goal

Let the expected time to the goal starting from (*n, n* − 1) and (*n* − 1*, n*) be denoted *T*_+_(*n*) and *T*_−_(*n*) respectively. We aim to compute *T*_+_(*N*) with an absorbing boundary at (1, 0), so that *T*_+_(1) = 0. The reflecting boundary conditions at (*N* − 1*, N*) imply *T*_−_(*N*) = *T*_+_(*N*) + 1. We have

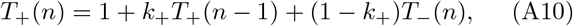

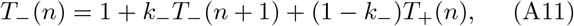

for 1 < *n* ≤ *N* and 1 < *n* < *N* respectively. Expressing *T*_−_(*n*) and *T*_−_(*n*+1) in terms of *T*_+_(*n*−1)*, T*_+_(*n*)*, T*_+_(*n*+1) using (A10) and plugging into (A11), we obtain the second-order difference equation

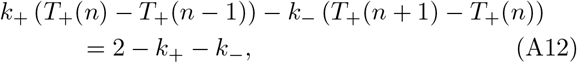

for 1 < *n* < *N*. Defining *U* (*n*) = *T*_+_(*n* + 1) − *T*_+_(*n*) gives

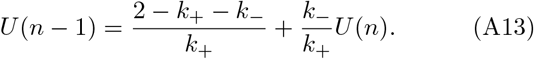

Since *T*_−_(*N*) = *T*_+_(*N*) + 1, setting *n* = *N* in (A10) gives *U* (*N* − 1) = −1 + 2*/k*_+_. Multiplying both sides by (*k*_−_*/k*_+_)^*n*−1^ and summing from *n* + 1 to *N* − 1, we get

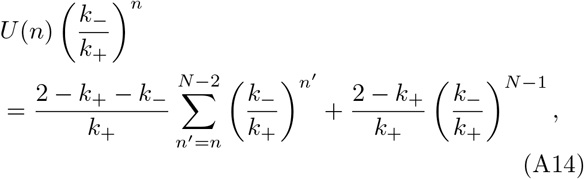

which after simplification leads to

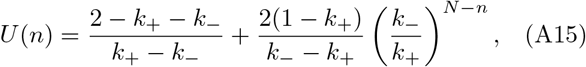

for *k*_−_ ≠ *k*_+_. Summing *U* (*n*) from *n* = 0 to *N* − 1 and using *T*_+_(0) = 0, we obtain the mean time to find the goal

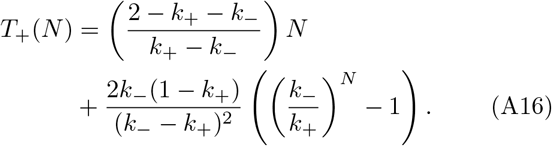

If *k*_−_ > *k*_+_, the agent takes time exponential in *N*, which is infeasible for large *N*. For *k*_−_ < *k*_+_, the agent travels linearly towards the goal with drift 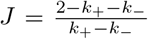. The marginal case of *k*_−_ = *k*_+_ leads to diffusive behavior. In this case, from (A14), we have

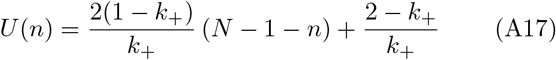

Summing from *n* = 0 to *N* − 1 gives 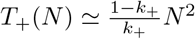.

## (b) Expected number of visits

We calculate the expected number of times per episode the agent visits each state on the direct path. Denote the expected number of times the agent visits states (*n* + 1*, n*) and (*n, n* + 1) as *M*_+_(*n*) and *M*_−_(*n*) respectively. We have

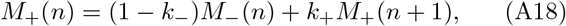

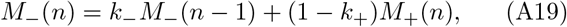

We observe that setting *n* → *n* + 1 in (A19) and adding (A18) gives a conservation equation, *M*_+_(*n*) + *M*_−_(*n* + 1) = *M*_−_(*n*) + *M*_+_(*n* + 1). At the boundaries we have *M*_−_(1) = (1−*k*_+_)*M*_+_(1) and *M*_+_(*N* −1) = *M*_−_(*N* −1)+1. Combining the latter boundary condition and the conservation equation, we see that *M*_+_(*n*) = *M*_−_(*n*) + 1 for all *n*. Plugging this into the former boundary condition then leads to *M*_+_(1) = 1*/k*_+_ and into (A18) leads to

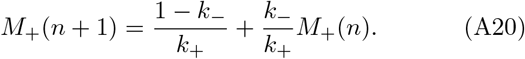

Dividing both sides by (*k*_−_/*k*_+_)^*n*+1^, summing from 1 to *n* and simplifying gives

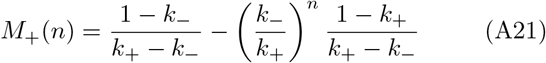

for *k*_−_ ≠ *k*_+_. When *k*_−_ = *k*_+_, we have *M*_+_(*n*) = 1 + *n*(1 − *k*_+_)*/k*_+_.

## Dynamics of learning

We now analyze the dynamics of reinforcement propagation for five learning rules, beginning with the SARSA rule considered in the main text. Throughout, we consider the slow-learning limit *α* ≪ 1. In this limit, we can compute the learning dynamics averaged over the agent’s behavior in each episode. Simulations show that the analytical results are accurate up to *α* ≲ 0.1, with wave-like propagation of the reinforcement signal observed for even larger *α*. In all cases analyzed below, we set the standard discount factor in RL (*γ*) to unity. This discount factor introduces an effective length scale for the influence of the reward and regularizes the Bellman equation, which are not necessary in our setting due to the regularization and effective horizon provided by the exploration-based stochastic policy. In our analysis below, it is easy to see that the effective horizon is *N*_range_ (−ln *σ*(*ε* + *r*))^−1^. For *N* ≪ *N*_range_, the reinforcement signal propagates as a wave. The wave slows down and the reinforcement signal eventually decays to zero at steady state when *N* ≫ *N*_range_. We consider the 1 ≪ *N* ≪ *N*_range_ limit in our analysis below, which thus requires that *ε* + *r* be appropriately large. In the linear track setting, reward is acquired only when the agent reaches the goal state. This is set by the boundary condition *q*_0_ = *r* (see the setup and notation section above). The initial conditions are 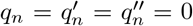 for *n* > 0. Also note that 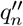, i.e., the *q_r_* value corresponding to going *away* from the goal is never reinforced and stays fixed at 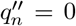. We thus consider the dynamics of *q_n_* and 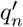 in what follows.

## (a) SARSA

Whenever the agent makes a state-action-state transition *s_t_, a_t_, s*_*t*+1_, the SARSA(0) (state-action-reward-state-action) learning rule updates the corresponding *q_r_* values as

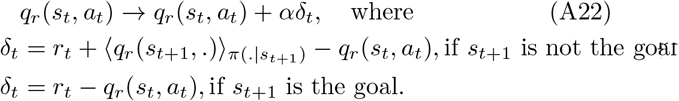

Here 0 < *α* < 1 is the learning rate, *r_t_* is the reward obtained at time *t* and the expectation is over future actions drawn from the policy *π*. As discussed previously, ln *π*(*a*|*s*) = *q_ε_*(*s, a*) + *q_r_*(*s, a*) − ln *Z*(*s*). Note that the learning rule described above is strictly speaking the “expected SARSA” rule [14], which is an efficient variant of the SARSA(0) rule. The 0 in the parenthesis corresponds to the eligibility trace parameter *λ* = 0, which converts the local SARSA(0) rule to a non-local rule for *λ* > 0. We discuss eligibility traces further below.

Applying the SARSA rule (A22), the expected change Δ*q_n_* and 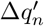 *in each episode* is given by

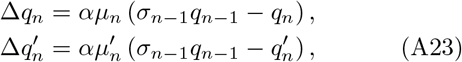

for *n* > 1 and Δ*q*_1_ = *αμ*_1_(*r* − *q*_1_), 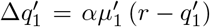. The *σ*_*n*−1_ *q*_*n*−1_ term is the expected value of *q_r_* at the subsequent state, i.e., once the agent crosses node *n*. Here *μ*_*n*_ and 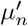 are the expected number of times per episode the agent crosses node *n* towards the goal along the direct path or by making a detour through the side path at *n*, respectively. The relative probability of these two events determines their ratio, 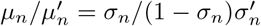, and the total expected crossings through node *n* towards the goal determines their sum 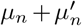. The expected crossings at each node depends in general on the *q_ε_* and *q_r_* values at all states. It is possible to calculate the expected crossings recursively similar to the case of pure exploration above. However, we note that for the nodes where learning occurs (near the front of the wave), we should expect a single crossing, i.e., 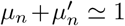. As argued in the main text, the key idea is that learning at node *n* only occurs when *σ*_*n*−1_ *q*_*n*−1_ is non-negligible, i.e., the probability of staying on the direct path is sufficiently reinforced at *n* − 1 and thus also for *n*′ ≤ *n* − 1. Since these actions are reinforced, the agent is very likely to take the direct path to the goal immediately after crossing *n*. Equivalently, whenever the agent crosses *n* towards the goal, the probability of it cycling back to *n*′ > *n* and making another attempt at crossing *n* is small, leading to 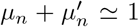. We thus obtain

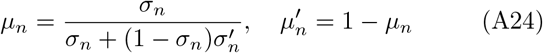

We can convert the discrete dynamics equations above to a continuous-time equation in the limit *α* ≪ 1, to get

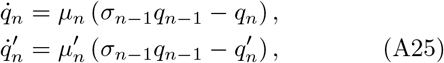

where the dot represents a time derivative and the unit of time is 1*/α*. Integrating the above equations over a unit time interval Δ*t* = 1 corresponds to expected changes in *q_n_* and 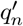 over 1*/α* episodes.

As shown in Figure 3a, integrating the equations (A25) provides an accurate approximation of the values obtained through full-scale RL simulations. We expand on the analysis of (A25) presented in the main text. Below, we consider exploration parameter values, *ε, ε*′*, ε*′′, such that *k*_−_ < *k*_+_, i.e., in the regime where the agent consistently gets to the goal and learning occurs. We set *ε*′′ = 0 and vary *ε, ε*′. The key parameter which determines the character of the wave is *ε*. *ε*′ is chosen such that *k*_−_ < *k*_+_. It is useful to consider the asymptotic limits *e^ε^* ≫ 1 (expanding regime) and *e*^−*ε*^ ≫ 1 (marching regime). The motivation for the names will become apparent.

## Expanding regime

The expanding regime has straightforward linear learning dynamics. To see this, note that when *e^ε^* ≫ 1, we have *σ_n_* ≃ 1 for all *n*. This implies from (A24) that *μ_n_* ≃ 1, 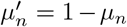 is negligible and 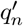 thus remains at 0. From (A25)

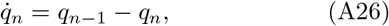

with boundary conditions *q*_0_(*t*) = *r* and initial conditions *q_n_*(0) = 0 for *n* > 0. Defining 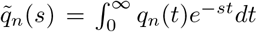, i.e., the Laplace transform of *q_n_*(*t*), (A26) leads to

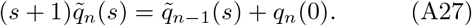

Multiplying both sides of the equation by (*s* + 1)^*n*−1^ and summing the recursion gives

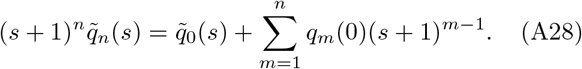

Using 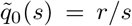 and performing the inverse Laplace transform leads to

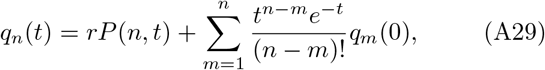

where *P* (*n, t*) is the regularized lower incomplete gamma function. Setting the boundary conditions, we have *q_n_*(*t*) = *rP* (*n, t*). For large *n* it is well known that the gamma distribution is approximated by a normal distribution: 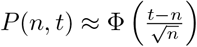 where Φ is the standard normal cdf. The half-maximum of the propagating signal thus travels as *n*_1/2_ = *t*, i.e., with speed *v*_0_ = 1 corresponding to 1 node every 1*/α* episodes. The width of the front expands as 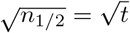.

## Marching regime

Qualitatively different behavior is observed in the opposite asymptotic limit, *e*^−*ε*^ ≫ 1. In this case, the nonlinear dynamics of (A25) come into play. We show below that the dynamics consist of a self-similar wavefront which marches forward one node at a time, with a fixed time interval, *τ*_0_, between each node (and thus its speed is 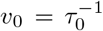). We first present the qualitative picture and then proceed to calculate *τ*_0_. It turns out that a complete characterization of the wave dynamics is feasible, which leads to analytical expressions for *τ*_0_, the shape of the wavefront and the dynamics of reinforcement at each node, *q_n_*(*t*).

Since *e*^−*ε*^ ≫ 1, the probability that the agent takes the direct path through node *n*, *μ_n_*, is negligible before any reinforcement occurs at node *n*. Any reinforcement that occurs is due to rare events which gradually increase *q_n_* provided that *q*_*n*−1_ > |*ε*|. This bottleneck remains until *q_n_* itself gradually increases to *q_n_* ≃ *ε*. Meanwhile, all the actions on the direct path for *n*′ < *n* are continuously reinforced. This reinforcement is possible as the agent can bypass crossing *n* through the direct path by instead taking a detour through the side path at *n*. Indeed, learning occurs (*k*_−_ < *k*_+_) in the marching regime only if *ε*′ is sufficiently large to allow for frequent detours through the side path. As a consequence, while *q_n_* is gradually being reinforced, 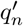 is sufficiently reinforced so that 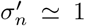 and *μ_n_* ≃ *σ_n_* (from (A24)). The detours through the side paths ensure that *q*_*n*−1_ is rapidly reinforced and *r* ≫ |*ε*| implies that *σ*_*n*−1_ displays switch-like behavior once *q*_*n*−1_ > |*ε*|. Since *q*_*n*−1_ is increasing, successive rare events through the direct path at node *n* receive increasing amounts of reinforcement on *q_n_*. Calculating the amount of reinforcement on *q_n_* thus requires knowing the dynamics of *q*_*n*−1_. However, the dynamics of *q*_*n*−1_ depend on the dynamics of *q*_*n*−2_, which in turn depends on *q*_*n*−3_ and so on, leading to cascading dependencies. The duration *τ*_0_ for *q_n_* to be reinforced from 0 to *ε* will thus depend on interactions with the bulk of the wave. We now calculate *τ*_0_ by solving the equation hierarchy. We will see that the solution is feasible due to the linear dynamics in the bulk of the wave (A26), which considerably simplify the analysis.

We begin (*t* = 0) from the moment when the wave has just reached node *n*, i.e., *q*_*n*−1_(0) = |*ε*| and *q_n_*(0) = 0 begins to be reinforced. *τ*_0_ is the time it takes for one cycle to complete, which is when *q_n_*(*τ*_0_) = |*ε*| is reached and *q_n_* begins to influence *q*_*n*+1_. From (A25), the (approximate) evolution of *q_n_* during this interval is given by

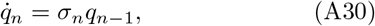

where we have used *μ_n_* ≃ *σ*_*n*_ and *σ*_*n*−1_ ≃ 1. Since *r* ≫ |*ε*| implies *q_n_* ≪ *q*_*n*−1_ in this period, we have also ignored the negative feedback due to *q_n_*. This argument can be made rigorous by computing *q_n_* from (A30) (see below), which will lead to an upper bound on *q_n_*, and showing that this upper bound is ≪ *q*_*n*−1_. Integrating (A30) for *t* ≤ *τ*_0_, we get

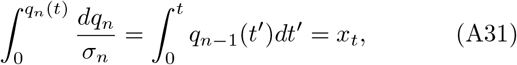

where 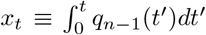 is to be calculated. Plugging in 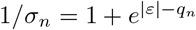 and integrating gives

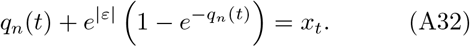

The solution of the above equation is expressed in terms of the Lambert *W* function [36], *W* (*x*),

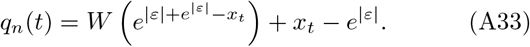

Since *q_n_*(*τ*_0_) = |*ε*|, *τ*_0_ is obtained from solving

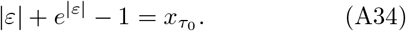

We now calculate *x_t_*. In the bulk (*m* < *n*), we have

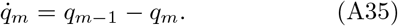

From (A29),

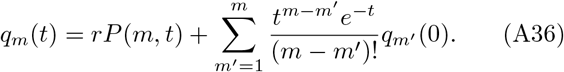

Note that the initial conditions *q_m′_* (0) here are the unknown values of *q_m′_* when *q*_*n*−1_ = |*ε*| and *q_n_* = 0, which are to be computed self-consistently under the marching dynamics with time step *τ*_0_. Using the series expansion of *P* (*m, t*) and *m*′ → *m* − *m*′, we re-write the above equation as

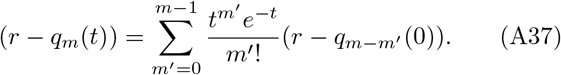

Here increasing values of *m*′ correspond to the nodes reinforced in earlier stages of the learning process. Thus, the terms *r* − *q*_*m*−*m*′_ (0) are decreasing in magnitude and the terms of large *m*′ do not matter for *τ*_0_ > 1. We may then take the sum to infinity in the equation above.

To solve for *q_m_*(0) for *m* < *n*, we notice that the self-similarity of the wavefront with period *τ*_0_ implies *q_m_*(*τ*_0_) = *q*_*m*−1_(0). Plugging *t* = *τ*_0_ into (A37), using *q_m_*(*τ*_0_) = *q*_*m*−1_(0), we get

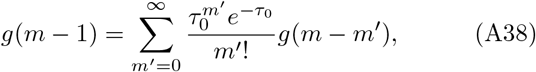

where *g*(*m*) ≡ 1 − *q_m_*(0)*/r*. This difference equation applies for all *m* < *n*. Since the difference equation has constant coefficients, solutions are of the form *g*(*m*) = *cb^m^*, where *c* is a constant [37]. We have

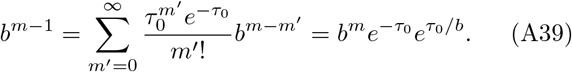

Thus *b* satisfies

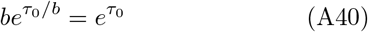

which leads to

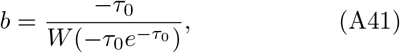

where *W* is the non-trivial branch of the Lambert *W* function (the trivial branch evaluates to 1). Evaluating (A37) for *m* = *n* − 1 then yields

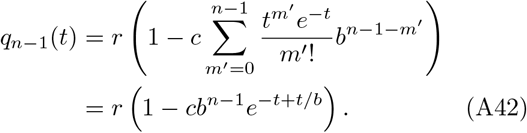

Since *q*_*n*−1_(0) = |*ε*|, we get *cb*^*n*−1^ = 1 − |*ε*|*/r*. The evolution of *q*_*n*−1_(*t*) is thus

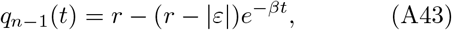

where 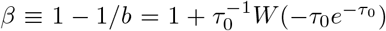, which determines the wave profile at any instant *t* since the periodicity implies *q*_*n*−*n*′_ (*t*) = *q*_*n*−1_(*t* + (*n*′ − 1)*τ*_0_).

Note that *W* (*x*) is real-valued for *x* ≥ − 1*/e*, which corresponds to *τ*_0_ ≥ 1. For *τ*_0_ = 1, using *W* (−1*/e*) = −1 we see that *β* = 1 and the power solution *g*(*m*) = *cβ^m^* cannot be used. It can be checked that *g*(*m*) = *cm* satisfies the recursion (*c* is again a constant). The initial conditions yield 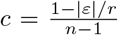, whose dependence on *n* violates the self-similarity of the wavefront. Thus, the dynamics when *τ*_0_ = 1 cannot be described by a self-similar traveling wave. Indeed, as shown in the expanding regime section, when *τ*_0_ = 1 the width of the wavefront expands as 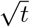.

Using the identity *e*^*W*(*x*)^ = *x/W* (*x*), we obtain 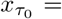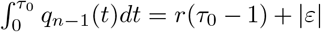. Plugging this into (A34), we finally have

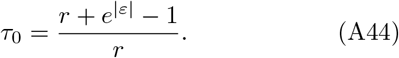

An interesting alternative derivation of *τ*_0_ exploits a conservation equation for 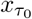. Since 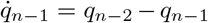, integrating both sides from 0 to *τ*_0_ and re-arranging yields

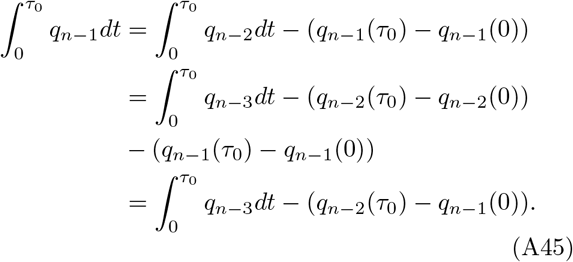

where we have used the periodicity in the bulk, *q*_*n*−2_(0) = *q*_*n*−1_(*τ*_0_) in the last step. Repeating this sequence of steps, we are lead to

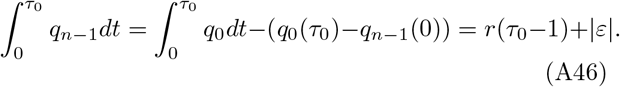

## (b) Eligibility traces

In RL, eligibility traces are used to enhance learning speed by efficiently propagating errors backwards in time. Specifically, the method uses the reward prediction error (*δ_t_* in (A22)) at the current state-action pair to update the *q_r_* values of recently visited state-action pairs, in addition to the *q_r_* values of the current state-action pair. The resultant *q_r_* update of the state-action pair visited *j* steps before the current one is

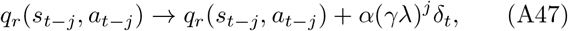

where 0 ≤ *λ* < 1 is the eligibility traces parameter and *γ* is the discount factor (recall that *γ* is set to 1 throughout our analysis). *λ* sets the effective number of previous state-action pairs which are affected by the update at the current state-action pair. We consider a slightly modified version of this learning rule, where *λ* is set to 1 and instead a fixed number *k* of previous state-action pairs are updated according to (A47).

Let us examine the effect of this rule when updating *q_n_* (the case of 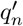 follows). If, after crossing node *n* towards the goal, the agent goes into the side path at *n* − 1, then since all the “hidden” state-action pairs inside the side path have *q_r_* = 0 and the side paths are assumed to be long detours (≫ *k*), the updates within the side path and after exiting the side path do not update *q_n_*. In this case, *q_n_* is updated just as in (A25). If the agent, instead of turning into the side path at *n* − 1, continues along the direct path towards node *n* − 2, and if *k* ≥ 1, then *q_n_* is updated as

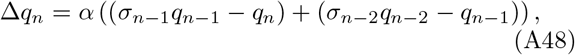

where the first term in the parenthesis is the *δ*-error from when the agent crosses node *n* and the second term is from when the agent crosses *n* − 1 along the direct path. If, at this point, the agent turns into the side path at *n* − 2, there are no further updates of *q_n_* for the same reason stated above. Similarly, if it instead continues along the direct path to *n* − 3 and *k* ≥ 2, then *q_n_* is again updated using the *δ*-error at that transition. Extending this argument, we see that *q_n_* (and 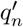 by the same argument) receives non-local updates as long as the agent continues along the direct path without taking turns into side paths. In the limit of *α* ≪ 1, we take expectations to get

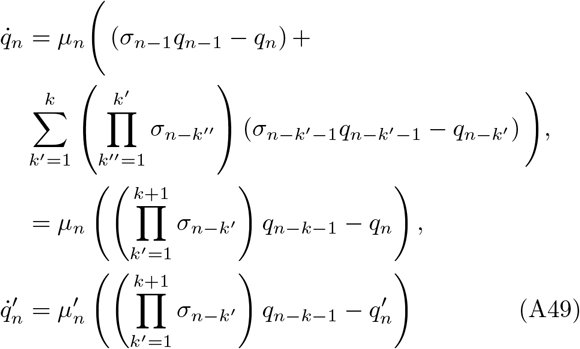

The product, 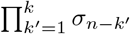, corresponds to the probability of taking *k* contiguous steps along the direct path. The second equation is obtained by noticing that all the terms except the ones involving *q_n_* and *q*_*n*−*k*−1_ cancel out. We verify the validity of (A49) by comparing to full RL simulations (Figure S3a).

As in the SARSA case, we consider the asymptotic limits *e^ε^* ≫ 1 and *e*^−*ε*^ ≫ 1. For *e^ε^* ≫ 1, we have *σ_n_* ≃ 1, *μ_n_* ≃ 1 for all *n* and the sums in (A49) collapse into the simple set of linear equations

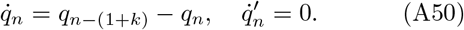

For convenience, we assume *n* = *ℓ*(1 + *k*) where *ℓ* is an integer. Similar results with minor modifications are obtained for the other cases. Taking the Laplace transform, we get

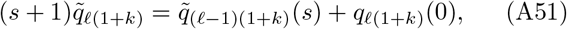

which after following steps similar to the SARSA case leads to

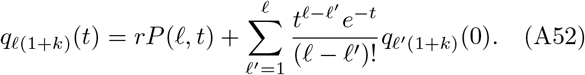

For the initial conditions *q_n_*(0) = 0 for *n* > 0, we have 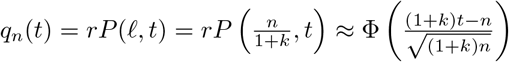 for large *n*. From here, we obtain the speed, *v_k_* = 1 + *k*, and the width of the wavefront at time *t*, 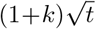. Thus, eligibility traces enhance the speed of reinforcement propagation by breaking the locality constraint of SARSA.

In the marching regime, *e*^−*ε*^ ≫ 1, suppose, as in the SARSA case, that the wave has just reached node *n*, i.e., *q*_*n*−1_(0) = |*ε*|, *q_n_*(0) = 0. We compute the time *τ_k_* it takes for *q_n_* to be reinforced to |*ε*|. When *q*_*n*−1_ ≥ |*ε*|, from (A41), the product of *σ*’s equals 1 and *q_n_*(*t*) subsequently evolves as

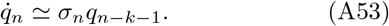

*τ_k_* can be calculated by integrating the above equation from *t* = 0 to *t* = *τ*, which gives

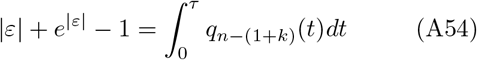

The integral 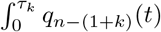 is determined by the dynamics in the bulk of the wave, which are governed by (A50). The Laplace transform of (A50) leads to a relationship similar to (A37) for *m* = *ℓ*(1 + *k*) < *n*:

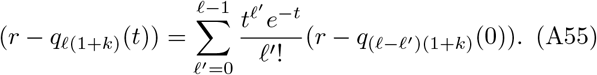

Defining *h*(*ℓ*) ≡ 1−*q*_*ℓ*(1+*k*)_(0)*/r* and using the periodicity of the wave w.r.t *τ_k_*, *q*_*ℓ*(1+*k*)_((1 + *k*)*τ*_*k*_) = *q*_(*ℓ*−1)(1+*k*)_(0), we have

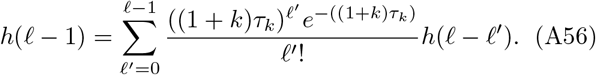

It can be checked that 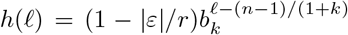with

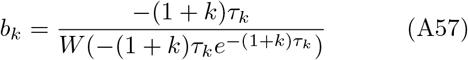

satisfies the above recursion and the boundary conditions *q*_*n*−1_(0) = |*ε*|. Defining *β_k_* ≡ 1 − 1*/b_k_*, we obtain

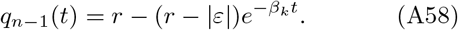

From (A54), we have

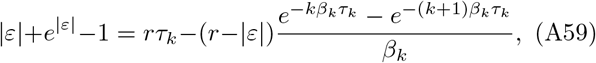

which upon rearranging leads to

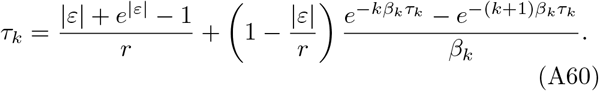

This implicit equation can be solved numerically for *τ_k_*. We verify the result by comparing our analytical calculation to RL simulations (Figure S3b,c). Importantly, it can be shown that the second term on the right-hand-side goes to zero as *k* → ∞. This limit corresponds to the situation in which the node *n* effectively receives reinforcement from future states that have already been fully reinforced to *r* (i.e., 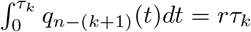). From here, we obtain the upper bound on the speed,

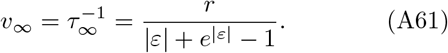

## (c) Dyna-Q

Dyna-Q (tabular) is a model-based RL algorithm which combines “planning” and *Q*-learning. It is useful to examine the learning dynamics of a model-based learning algorithm when applied to our setting. The Dyna-Q agent learns a model of the environment, *P* (*s*′*, r|s, a*), corresponding to the distribution of subsequent rewards and states for every state-action pair. At every step during navigation, in addition to updating its *q*-values using (A63) for the current state-action pair, the agent also randomly samples *n_p_* previously visited state-action pairs from memory, draws the subsequent state and reward from its learned model, and updates the corresponding *q*-values.

Dyna-Q does not significantly influence the learning dynamics in our setting due to the deterministic state transitions and rewards. Its effect is to simply scale the learning rate compared to SARSA. This scaling is because the probability, *p*(*s, a*), that a particular state-action pair, (*s, a*), is reinforced during each of the additional *n_p_* planning steps is proportional to the number of times (*s, a*) is expected to be traversed in an episode, say, *n*(*s, a*). The number of times (*s, a*) is reinforced in a single planning step is then 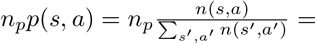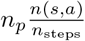, where *n*_steps_ is the total number of state-action pairs visited in each episode. Since this planning step is applied at each step, the total *additional* number of times, *q_r_*(*s, a*) is updated per episode is *n_p_n*(*s, a*), in addition to the *n*(*s, a*) regular SARSA updates. This calculation implies that (*s, a*) is updated by a factor *n_p_* + 1 compared to the plain SARSA case, which can be interpreted as a scaling of the learning rate *α*→ (1 + *n_p_*)*α*. Here, we have assumed that the memory of the Dyna-Q agent is not much larger than *α*^−1^ so that *n*(*s, a*) does not change significantly over a timescale comparable to the memory size. We confirm this prediction in RL simulations of Dyna-Q agents (Figure S4a).

## (d) Q-learning

Watkins’ *Q*-learning rule is closely related to SARSA with an important difference that enables off-policy learning, i.e., the agent learns the optimal *q* values while executing an arbitrary policy. This rule is defined by

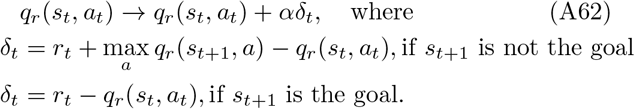

Note the max instead of the expectation in *δ_t_*. We consider *Q*-learning using the stochastic policy considered for SARSA above. The equivalent of (A25) follows by replacing *σ*_*n*−1_*q*_*n*−1_ in (A25) with max(0, *q*_*n*−1_) = *q*_*n*−1_ since the alternative action of turning into the side path has *q_r_* = 0 and *q*_*n*−1_ ≥ 0. We thus have

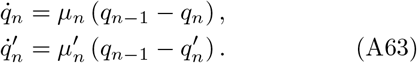

The *σ*_*n*−1_ prefactor does not play a significant role, and the results obtained for SARSA are directly applicable in both the expanding and marching regimes, which we verify through RL simulations (Figure S4b).

## (e) Q-learning (pure exploration)

Next, we consider *Q*-learning with a purely explorative policy that is independent of the learned *q_r_* values. Specifically, the policy is fixed and is given by ln *π*(*a|s*) = *q_ε_*(*s, a*) − ln *Z*(*s*), where *q_ε_*(*s, a*) are the intrinsic rewards as in the previous cases. This case is useful to understand the effects of the learning rule in isolation, without the effects of the feedback on behavior due to learning coming into play.

The dynamics of *q_n_* and 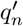 are given by (A63). However, the factors *μ_n_* are no longer the ones in (A24). Recall that *μ_n_* is the expected number of times per episode the agent performs the action of traversing towards the goal at node *n* on the direct path. This number is the expected visits to state (*n* + 1*, n*), *M*_+_(*n*) (from (A21)), multiplied by the probability, *σ*(*ε*), that the agent takes the action leading to (*n, n* − 1). We thus have *μ_n_* = *σ*(*ε*)*M*_+_(*n*) and, similarly, 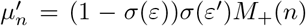. Since *μ_n_*’s are constant over time, the differential equations (A63) are linear and can be exactly solved for *q_n_*(*t*) in the Laplace domain. We obtain

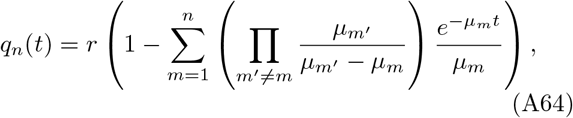

which unfortunately does not yield much insight. Instead, we consider the evolution of *q_n_*(*t*) for *n* large when *k*_−_ < *k*_+_. From (A21), we have that 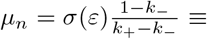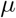 is a constant. The equations 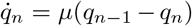 lead to *q_n_*(*t*) ≈ *rP*(*n*, *μt*). This relation is approximate as *μ_n_* is not constant for small *n*. The reinforcement signal thus propagates with speed

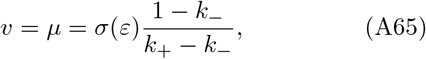

which is verified in simulations (Figure S4c). This analysis further highlights that when learning is decoupled from behavior, similar to the *e^ε^* ≫ 1 expanding regime, the propagation of the signal is simply constrained by the local learning rule with speed proportional to the number of times the learning rule is applied at each state-action pair.

---

**FIG. S1.**
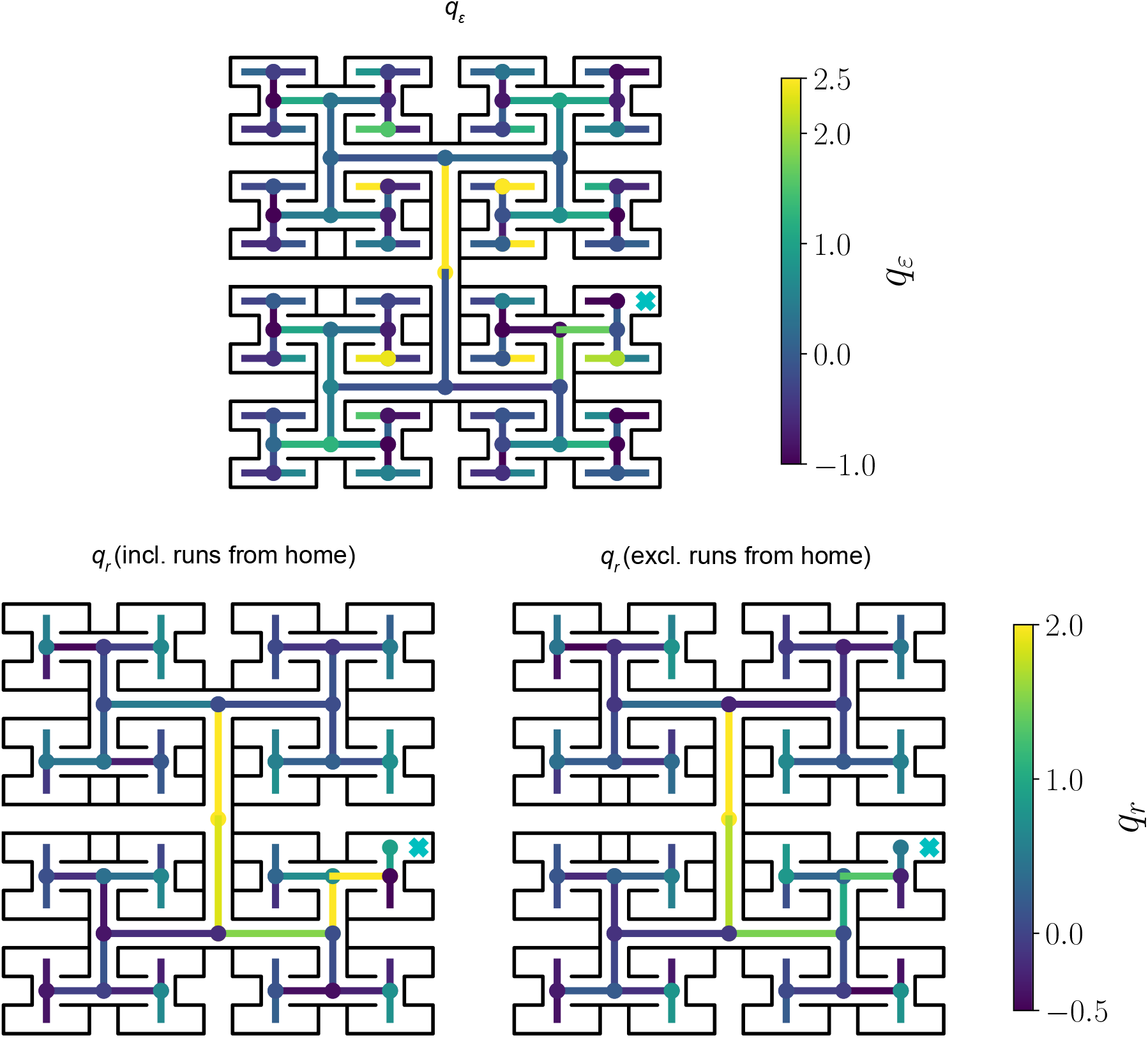
We estimate the *q_r_* values from experimental data using MaxEnt IRL by comparing the behavior of rewarded and unrewarded mice. Shown here are a subset of the *q_r_* values for the state-action pairs leading to the reward (blue cross). For example, the color of the edge immediately after taking a right turn from home represents the *q_r_* value of taking a left turn at the next junction. To ensure habitual direct runs from home to goal do not significantly bias the estimated *q_r_* values close to the reward, we show the estimates from data when runs from home are included (left) and excluded (right).

**FIG. S2.**
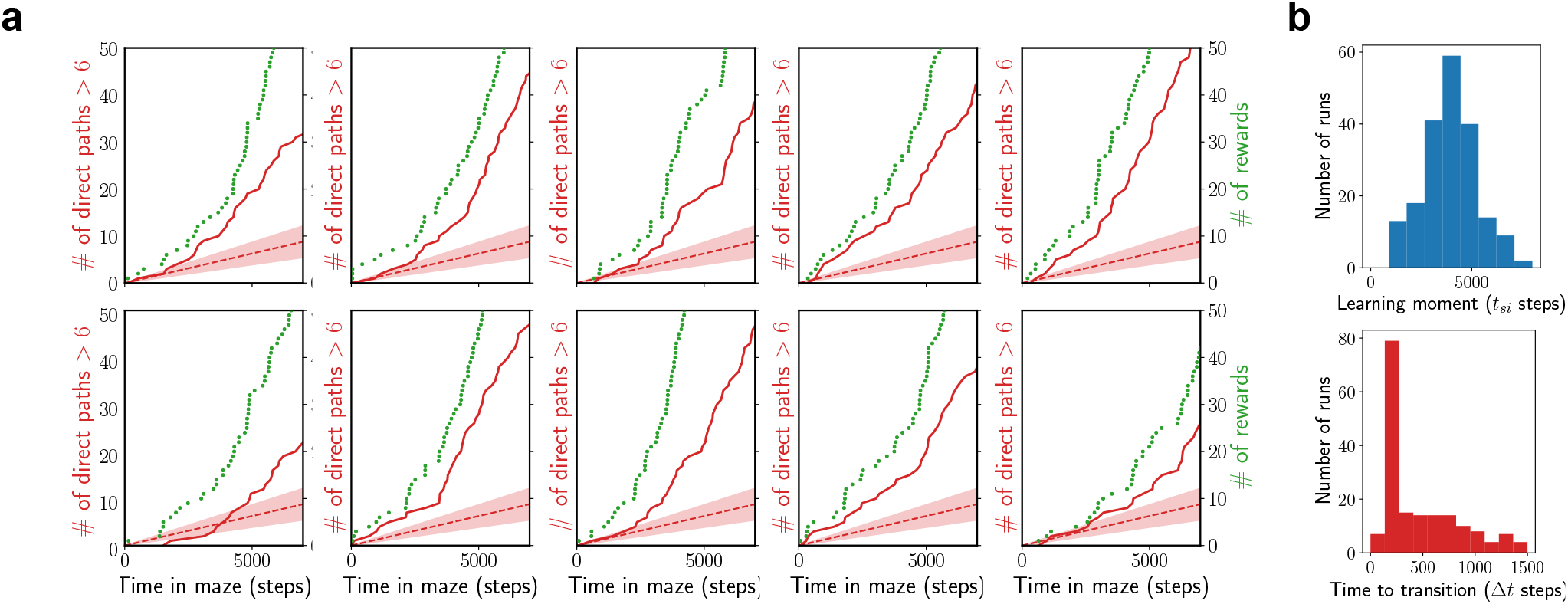
(a) Learning curves from ten random runs of the RL simulations. The dashed lines and solid fill correspond to the average ± one standard deviation in the rate of direct paths for unrewarded agents. (b) To quantify the variability across realizations, we repeat the RL simulations for 200 runs and fit the rate of the direct path for each run to a logistic function, 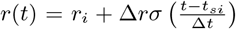 using the maximum likelihood method described in ref. [20]. The histograms for the best fit *t_si_* and Δ*t* are shown. Note the sharp peak at Δ*t* ≈ 150 steps (≈ 3 episodes).

**FIG. S3.**
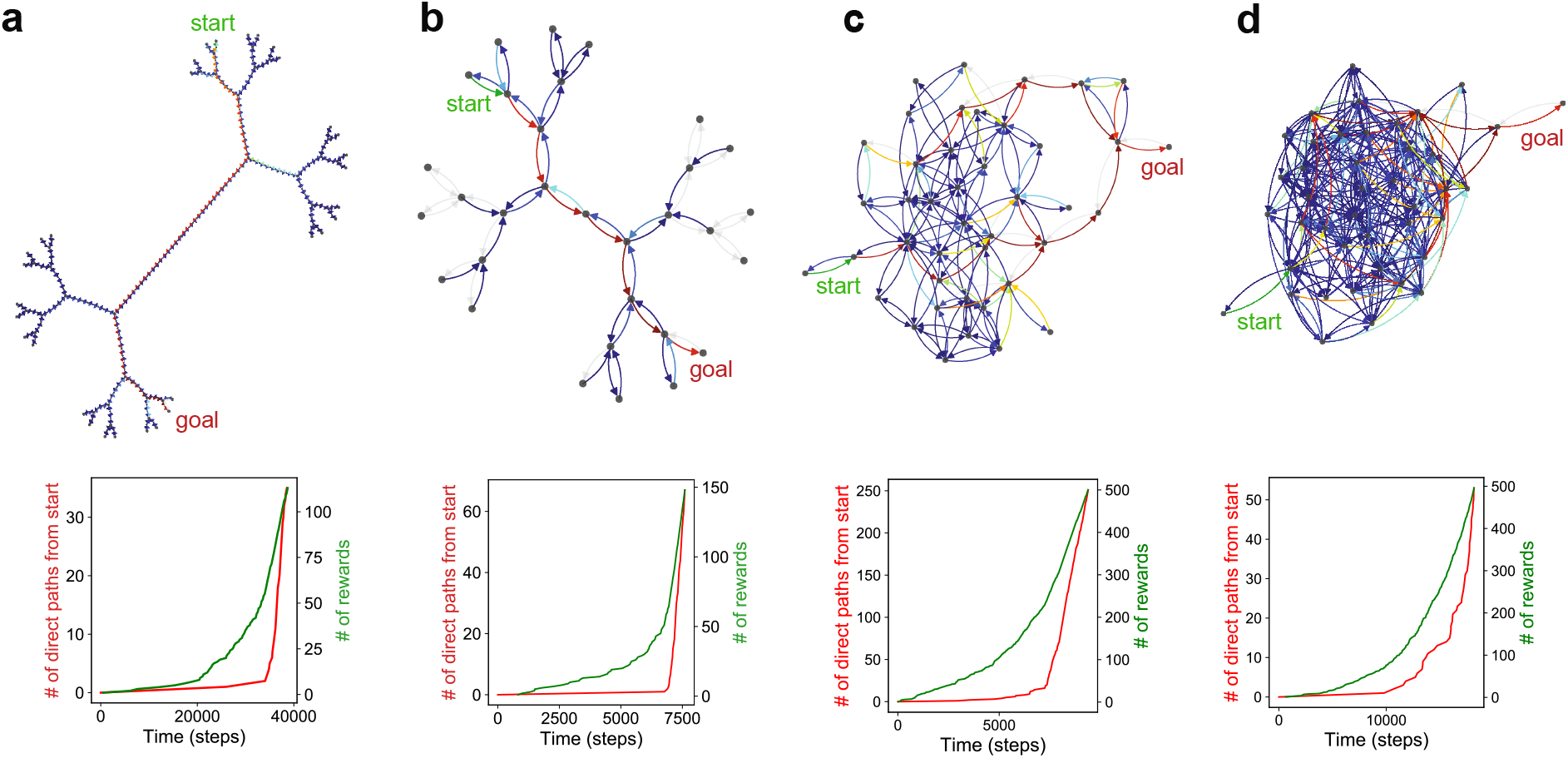
Numerics and theory for RL with eligibility traces. (a) The 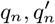 from RL simulations align closely with numerics (A49). (b) The speed from RL simulations and theory for a range of values of *k, ε, r*. (c) The shape of the wave front from numerics matches the theory prediction (A58). Here *k* = 2, *ε* = −2, *r* = 15.

**FIG. S4.**
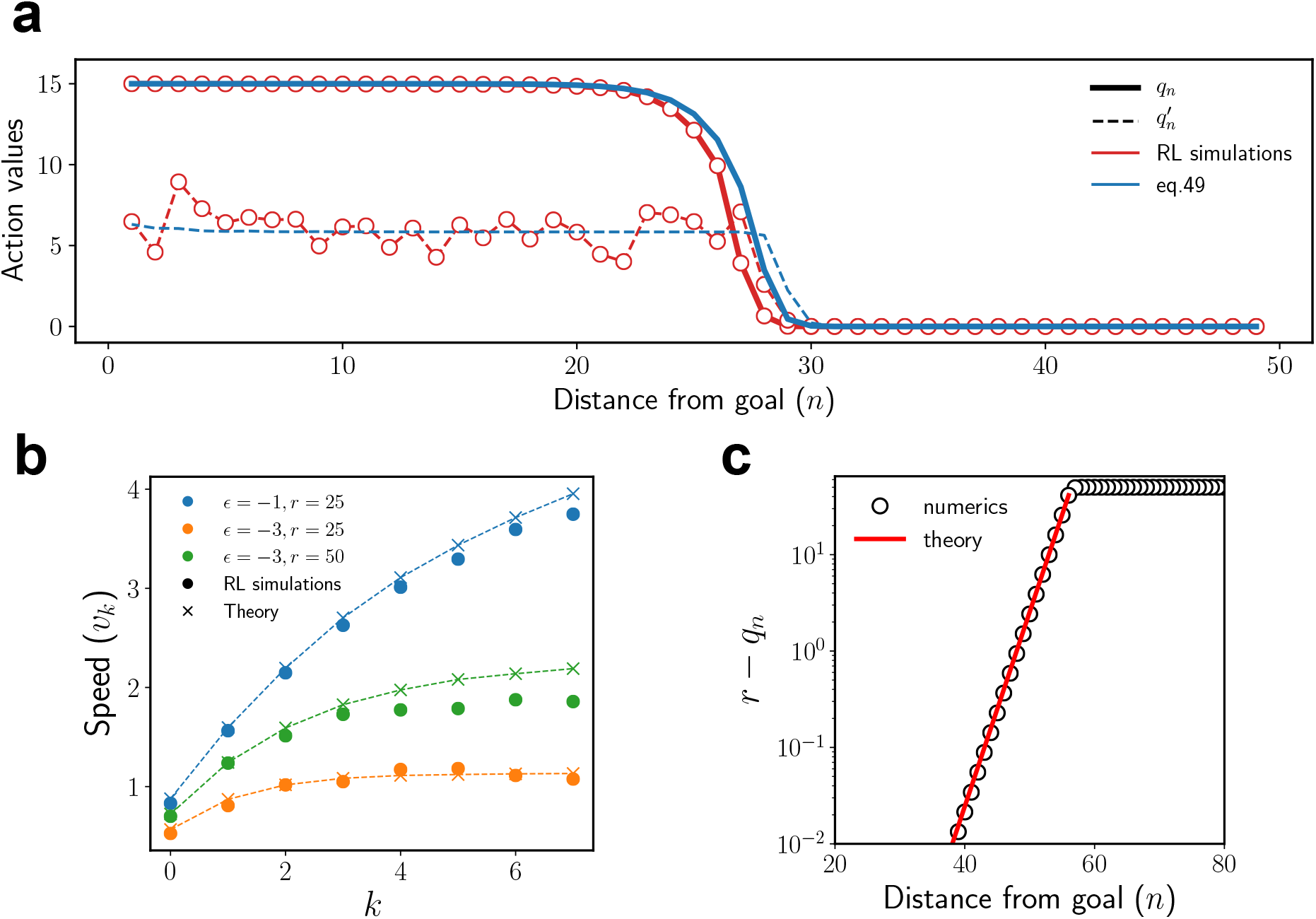
Theoretical predictions of the speed for various RL rules compared to those obtained from simulations.

## Notes

### Competing Interest Statement

The authors have declared no competing interest.

### Summary of Updates

The title, abstract and text have been re-framed to clarify the main message; New simulations have been added to highlight the generality of the result; Movies S2-S5 added; Figure 2 split with additional data into Figures 2 and 3; Discussion updated; Minor changes throughout the text for clarity

